# Human adolescent brain similarity development is different for paralimbic versus neocortical zones

**DOI:** 10.1101/2023.09.17.558126

**Authors:** Lena Dorfschmidt, František Váša, Simon R. White, Rafael Romero-García, Manfred G. Kitzbichler, Aaron Alexander-Bloch, Matthew Cieslak, Kahini Mehta, Theodore D. Satterthwaite, the NSPN consortium, Richard A.I. Bethlehem, Jakob Seidlitz, Petra E. Vértes, Edward T. Bullmore

## Abstract

Adolescent development of human brain structural and functional networks is increasingly recognised as fundamental to emergence of typical and atypical adult cognitive and emotional processes. We analysed multimodal magnetic resonance imaging (MRI) data collected from N ∼ 300 healthy adolescents (51%; female; 14-26 years) each scanned repeatedly in an accelerated longitudinal design, to provide an analyzable dataset of 469 structural scans and 448 functional MRI scans. We estimated the morphometric similarity between each possible pair of 358 cortical areas on a feature vector comprising six macro- and micro-structural MRI metrics, resulting in a morphometric similarity network (MSN) for each scan. Over the course of adolescence, we found that morphometric similarity increased in paralimbic cortical areas, e.g., insula and cingulate cortex, but generally decreased in neocortical areas; and these results were replicated in an independent developmental MRI cohort (N ∼ 304). Increasing hubness of paralimbic nodes in MSNs was associated with increased strength of coupling between their morphometric similarity and functional connectivity. Decreasing hubness of neocortical nodes in MSNs was associated with reduced strength of structure-function coupling and increasingly diverse functional connections in the corresponding fMRI networks. Neocortical areas became more structurally differentiated and more functionally integrative in a metabolically expensive process linked to cortical thinning and myelination; whereas paralimbic areas specialised for affective and interoceptive functions became less differentiated, as hypothetically predicted by a developmental transition from peri-allocortical to pro-isocortical organization of cortex. Cytoarchitectonically distinct zones of human cortex undergo distinct neurodevelopmental programmes during typical adolescence.

---

Magnetic resonance imaging (MRI) studies of human brain structure during adolescence and childhood have already identified two major developmental processes that are on-going during the maturational transition from birth to adult brain organization: (i) after peaking in early childhood, cortical grey matter volume and thickness monotonically decrease during adolescence; while (ii) protracted myelination of the cortex sees peak white matter volumes reached in early adulthood (Bethlehem et al. 2022). Anatomical MRI maps of cortical thinning and myelination markers like magnetization transfer (MT) are highly (negatively) correlated, indicating that these may be technically and/or biologically confounded measurements of the same underlying process of synaptic pruning and consolidation (Whitaker et al. 2016; Tamnes et al. 2017). These and other large-scale, long-term neurodevelopmental programs are thought to be fundamental to the emergence of adult cognitive functions and social behaviours (Váša, Seidlitz, et al. 2018; Mills, Goddings, Herting, et al. 2016; Sowell et al. 2004).

To date, neurodevelopmental MRI data have largely been studied using models of change in brain structure measured globally, or one region at a time (Mills, Goddings, Herting, et al. 2016; Mills, Goddings, Clasen, et al. 2014; Raznahan et al. 2011). It is now timely to understand more about developmental change in brain structure measured by network or connectomic metrics, given the dominant known adolescent process of pruning and consolidation of (synaptic) connectivity between neurons and regions. A technical challenge for developmental network neuroscience has been the estimation of anatomical or structural connectivity between hundreds of regional (cortical and subcortical) nodes in each individual’s brain scan. Previous studies have used structural covariance analysis of a single MRI metric measured at each region in a large group of scans (Váša, Seidlitz, et al. 2018; Whitaker et al. 2016); or tractography algorithms to reconstruct the fascicles and tracts of white matter projections between cortical areas (Baum et al. 2020). However, both approaches are problematic in different ways. Structural covariance analysis does not resolve the network organization of an individual scan, which is obviously crucial for within-subject developmental modeling of repeated measures in a longitudinal design. Tractography networks can be reconstructed from an individual’s diffusion weighted imaging (DWI) data; but the resulting connectomes generally suffer from under-estimation of long-range connections, e.g., between bilaterally symmetric cortical areas (Dauguet et al. 2007; Donahue et al. 2016; Maier-Hein et al. 2017), and have been unfavourably benchmarked against structural covariance and newer methods for estimating the anatomical connectome in a single scan (Seidlitz, Váša, et al. 2018a; Sebenius et al. 2023).

An alternative approach has emerged recently based on the concept that cytoarchitectonically similar regions are more likely to be axonally connected, as articulated in more detail by the structural model of Barbas and colleagues (Beul et al. 2017; García-Cabezas et al. 2019). On this basis, estimating the similarity between cortical or subcortical areas, each structurally phenotyped by one or more MRI metrics, could provide a tractable and plausible proxy for their axonal inter-connectivity (Haber et al. 2022). This principle of homophily - “like connects with like” - has been demonstrated in nervous systems across scales and species. At the microscopic (cellular) level, synaptic connections are more likely to form between two functionally and developmentally similar neurons; at the macroscopic (whole brain) level, animal models have shown that there is greater inter-areal connectivity via large-scale axonal tracts between regions that have similar laminar structure and cellular composition (Goulas et al. 2017). Lastly, generative modelling approaches have shown that a simple two parameter model of distance-penalised homophily (Vértes et al. 2012; Betzel et al. 2016) simulates many aspects of human brain network organization as an economic trade-off between wiring cost and topological complexity (Akarca et al. 2021).

Morphometric similarity translates this invasive or in silico work to in-vivo human brain networks by estimating all pair-wise inter-regional correlations of a multi-modal MRI feature vector measured at each region. MSNs can be built from any combination of structural MRI metrics (**Fig. 1A**), including (i) macro-structural metrics, like cortical thickness (CT), grey matter volume (GM) and surface area (SA), which aggregate data from multiple voxels representing an anatomical region to estimate its geometric properties on the centimetre scale; and (ii) micro-structural metrics, which are representative of some aspect of brain tissue composition on the millimetre scale of a single voxel, e.g., MT is widely regarded as a proxy for cortical myelination (**Fig. 1B,C**). MSNs have been shown to correlate with the “gold standard” of anatomical connectivity, axonal tract tracing data, in the macaque monkey, thus validating morphometric similarity as a proxy for axonal connectivity. Network phenotypes, like the degree of connectivity or “hubness” of each regional node in the connectome, have also been used to predict individual differences in intelligence (Seidlitz, Váša, et al. 2018b), to track normative brain network development in the first decade of life (using longitudinal data from the ABCD cohort) (Wu et al. 2022), and to discover case-control differences in brain network phenotypes across a range of rare genetic and other neurodevelopmental (Seidlitz, Nadig, et al. 2020), psychiatric (Morgan et al. 2019; Li, Seidlitz, et al. 2021), and neurological disorders (Li, Keller, et al. 2022).

**Fig. 1:**
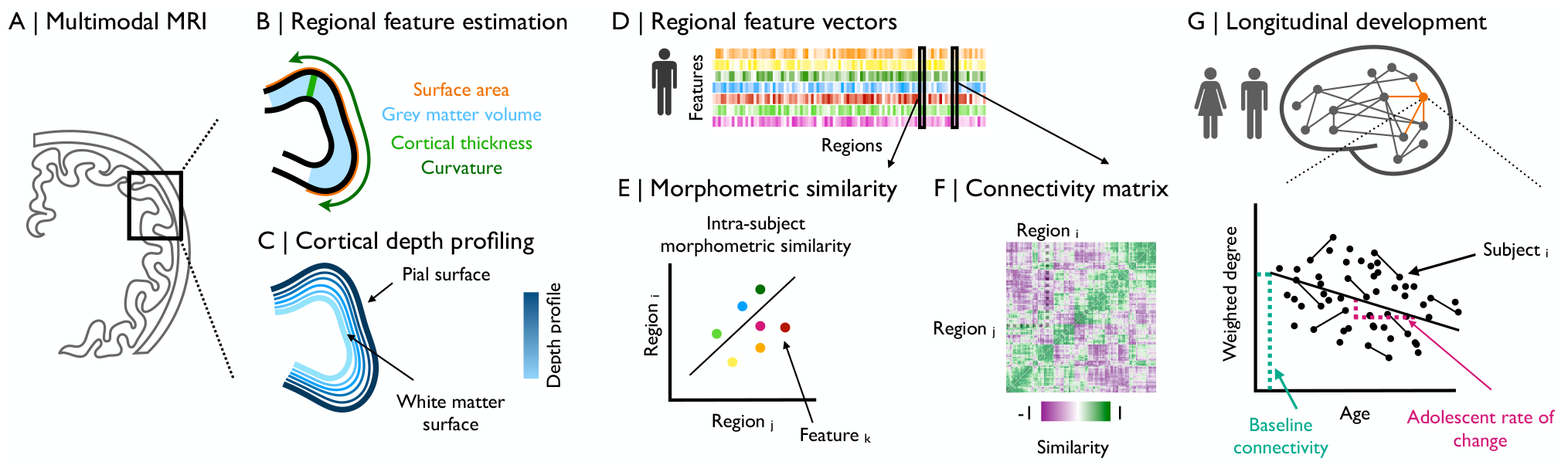
Modelling of adolescent changes in morphometric similarity: We estimated morphometric similarity networks (MSNs) from multi-modal MRI brain scanning data. (A) These images were parcellated into 358 pre-defined cortical regions. (B) Macro-structural MRI phenotypes, like CT, GM, and SA, were estimated for each cortical area overall. (C) Additionally, depth-dependent profiling was used to construct multiple cortical surfaces between the white matter surface and the pial surface for estimation of MT, a micro-stuctural MRI phenotype, with fine-grained laminar resolution at 70% of cortical depth. (D) We constructed a feature matrix of multiple features estimated at each region for each individual subject, resulting in a subject-specific *{Regions × Features}* multi-modal MRI data matrix. (E) We estimated the similarity between each pair of cortical areas in terms of the pairwise correlations between regional feature vectors comprising multiple normalized macro- and micro-structural MRI features estimated at each region. (F) We compiled all possible inter-areal similarity measures in a subject-specific *{Regions × Regions}*association matrix or morphometric similarity network. (G) We estimated adolescence-related changes in weighted degree, i.e., the mean weight over all of each node’s edges: the baseline functional connectivity, as the predicted nodal degree at age 14; and the rate of change of hubness, as the slope of a linear regression of age on weighted degree.

Here, we used MSNs to integrate multiple macro- and micro-structural MRI phenotypes, including features like cortical thickness and MT that have previously been linked separately to adolescent brain structural maturation (Whitaker et al. 2016), to measure developmental changes in individually estimated brain networks. We analysed structural MRI scans in an accelerated longitudinal design (N=291, age range 14 to 26 years; 51% female), stratified by age and balanced for sex per age stratum, with each participant scanned at least twice at 6-18 month intervals (**SI Fig. S1**). From the six macro- and micro-structural MRI metrics measured at each of 358 cortical areas, we estimated the morphometric similarity network for each of 469 individual scans. We used linear mixed effects models to estimate developmental change in MSN edge weights and nodal degree (a measure of hubness), and we compared these structural network results to functional networks constructed from 448 functional MRI scans in a large subset (N=283) of the NSPN sample. Recognising the importance of replicability, we endeavoured to reproduce key results from the NSPN cohort in an independent dataset, using N=304 cross-sectional scans from subjects 14-21 years collected as part of the Human Connectome Project Development sample (HCP-D).

We hypothesised (i) that there would be developmental changes in MSN phenotypes, e.g., some regional nodes might be become more or less hub-like over the course of adolescence, and (ii) that cytoarchitectonically distinct zones of cortex, e.g., paralimbic cortex compared to neocortex or isocortex, might have different trajectories of MSN development. We also predicted (ii) that adolescent changes in structural networks should be related to concomitant changes in functional network organization.

## Results

### >Analyzable sample

The final sample of morphometric feature data from the NSPN cohort after quality control consisted of 469 scans from 291 subjects in 358 regions. The fMRI sample included 448 scans from 283 subjects at 330 regions (**SI Table S1**). We conducted each analysis on the largest possible dataset, thus analyses of brain structure were conducted on 291 subjects across 358 regions, whereas analyses of structure-function relationships were conducted on a subsample of 283 subjects across 330 regions (**SI Table S2**). After quality control and age-matching, replication analyses were conducted using structural MRI data from N=304 subjects of the HCP-D cohort.

### >Adolescent changes in global and regional MRI metrics

We first estimated adolescent changes at global and regional scales for six morphometric features: (i) three macro-structural MRI metrics: cortical thickness, grey matter volume, and surface area; (ii) and three microstructural metrics: magnetization transfer, fractional anisotropy and mean diffusivity. We estimated the effect of age on these features using linear mixed effects (LME) models with a fixed effect of age, sex, and site, and a random effect of subject (**Fig. 1G**).

Globally, we found that all macro-structural metrics significantly decreased during adolescence: SA, *t*_*age*_ = −2.33, *P*_*FDR*_ *<* 0.05; GM, *t*_*age*_ = −5.23, *P*_*FDR*_ *<* 0.01); and CT, *t*_*age*_ = −7.29, *P*_*FDR*_ *<* 0.01. Of the microstructural metrics, MT significantly increased (*t*_*age*_ = 3.19; *P*_*FDR*_ *<* 0.01) while FA (*t*_*age*_ = 1.78) and MD (*t*_*age*_ = −0.42) showed no significant changes after correction for multiple comparisons (**Fig. 2A, SI Table 3**). We also found that there were significant sex differences in one feature, GM, *t*_*sex*_ = 2.85, *P*_*FDR*_ *<* 0.05. For the subset of subjects with a baseline and one year follow up scan, we further confirmed that in all features but FA these global trends are also observed in the raw within-subject differences in global feature strength between baseline and follow up (for details see **SI Fig. 4**).

**Fig. 2:**
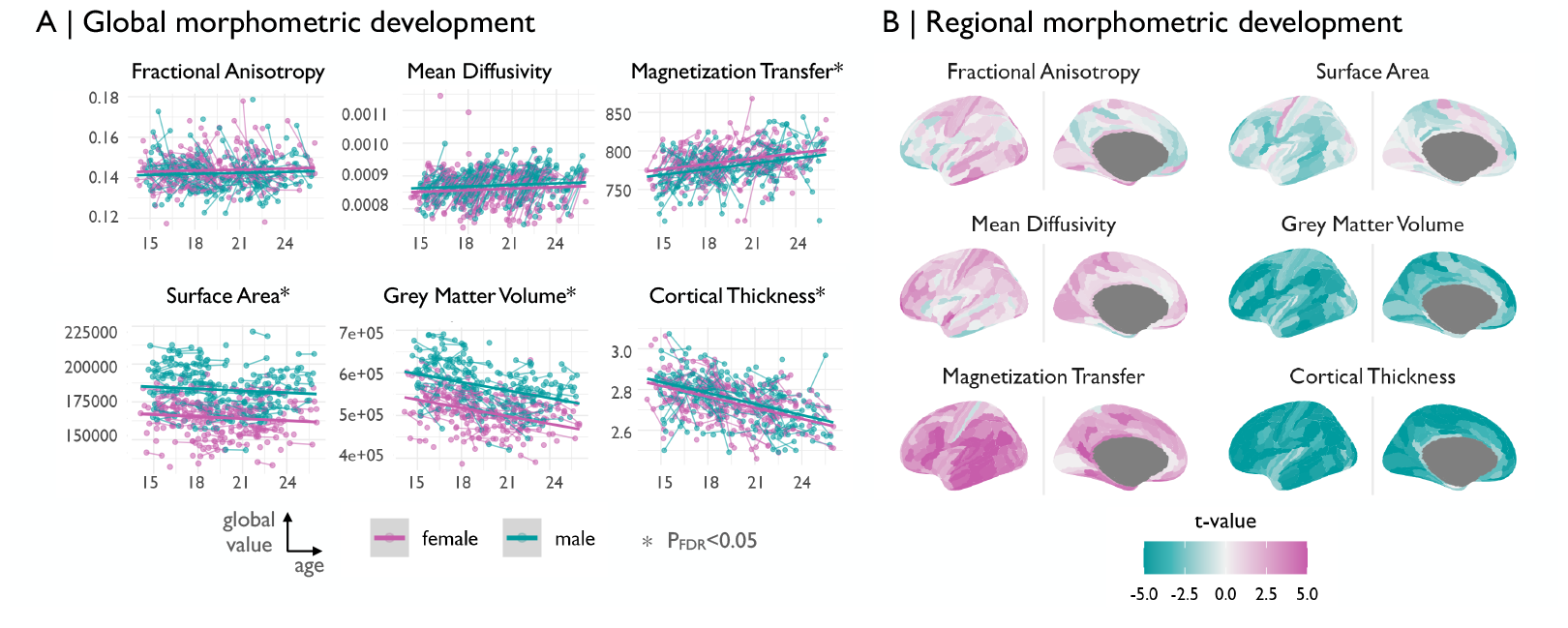
Adolescent changes in regional macro-structural and micro-structural MRI metrics: We modeled the linear effect of age on six morphometric features. (A) Globally, the three macro-structural MRI metrics (GM, CT, SA), all decreased over the course of adolescence (*P*_*FDR*_ *<* 0.05 for each), while one of the three micro-structural MRI metrics was significantly increased during adolescence (MT, *P*_*FDR*_ *<* 0.05), but not MD or FA. (B) We modeled the linear effect of age on six morphometric features at each of 358 cortical areas to resolve the anatomical patterning of developmental changes in macro- and micro-structural MRI metrics during adolescence. We generally observed increases (*t >* 0) in micro-structural features, and decreases (*t <* 0) in macro-structural features.

Mirroring the global developments, regionally, we found that macro-structural MRI metrics tended to decrease, and micro-structural metrics tended to increase (**Fig. 2B, SI Fig. S5**); We found that this effect was strongest for MT, where 289 regions significantly (*P*_*FDR*_ *<* 0.05) increased in weighted degree over the course of adolescence, and in CT and GM, where 336 and 296 regions, respectively, significantly decreased (for a map of change relative to a feature’s global development see **SI Fig. S6**). We found some variability between cytoarchitectonic zones of cortex in terms of their age-related changes in each of the MRI metrics, e.g., all cortical zones had decreased macro-structural metrics but the magnitude of shrinkage was consistently less in paramlimbic cortex compared to neocortical zones (**SI Fig. S7, SI Table S4**).

### Adolescent change in morphometric similarity

We constructed MSNs from each participant’s set of T1 and DWI MRI scans, at each time-point, by estimating the Pearson correlation between all pairwise regional feature vectors comprising the six MRI metrics, resulting in a *{*358 *×* 358*}* symmetric morphometric similarity matrix or weighted, undirected MSN. The weighted degree, *k*, of each regional node in each MSN is a measure of its morphometric similarity with all other regions, and high degree nodes or hubs are morphometrically similar to many other nodes in the brain.

Because we constructed a MSN model of the connectome for each scanning session completed by each participant, we could estimate developmental changes in MSN parameters using the same LME as previously used for analysis of age-related change in global and regional MRI metrics (**Fig. 1G**). We found that weighted degree generally decreased with age, i.e., Δ*k <* 0, in heteromodal and other neocortical zones, meaning that these areas became more morphometrically dis-similar from the rest of the brain; whereas degree increased with age, i.e., Δ*k <* 0, in paralimbic areas such as the insula and cingulate cortex, meaning they became more morphometrically similar to the rest of the brain (**Fig. 3A**).

**Fig. 3:**
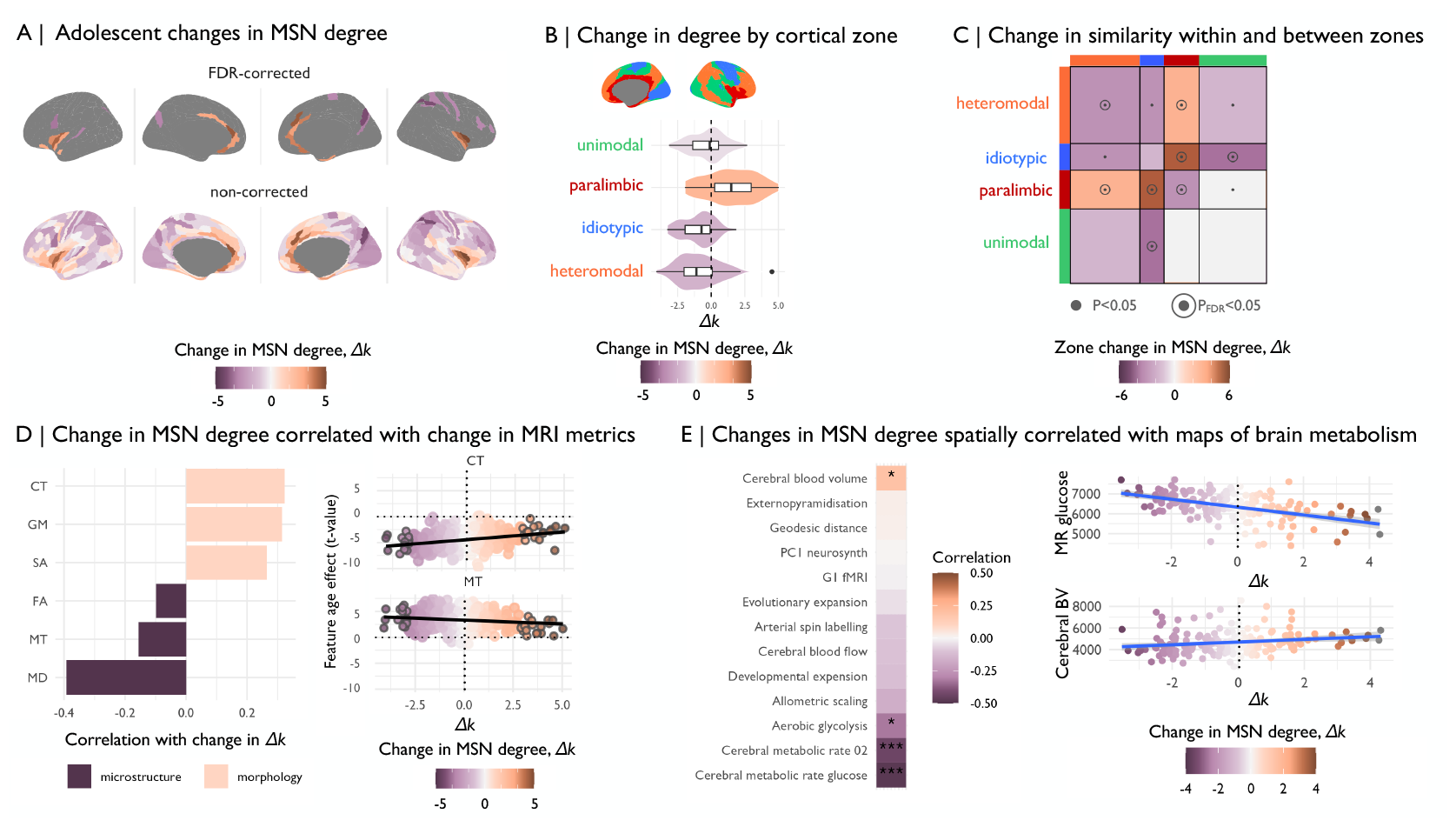
Adolescent change in degree of morphometric similarity, Δ*k*: We estimated morphometric similarity networks for each subject by correlating the standardized morphometric feature vectors for each possible pair of regions. (A) We estimated linear changes in morphometric similarity with age, Δ*k*, at each region and found that morphometric similarity decreased in neocortical (frontal, occipital) regions, and increased in medial and temporal cortical regions. These changes were significant after correction for multiple comparisons in 33 regions. (B) We estimated the mean effect of age on all regions within each of the Mesulam cytoarchitectonic zones (Mesulam 1998; Mesulam 2000) and found that morphometric similarity increased in the paralimbic cortex and decreased in all other zones. (C) We also found that within-network similarity decreased for all zones, marginally in unimodal and idiotypic zones and significantly after correction for multiple comparisons in heteromodal and paralimbic zones (*P*_*FDR*_ *<* 0.05), while between-network connectivity increased between paralimbic and hetermodal, as well as idiotyic zones (*P*_*FDR*_ *<* 0.05) and decreased otherwise. (D) Next, we assessed the correlation between adolescent effects on individual MRI features at each region and the adolescent effect on degree of morphometric similarity, or “hubness”, of each regional node in the cortical connectome. We found that MT and other micro-structural MRI features were negatively correlated with adolescent change in MSN degree, i.e., cortical myelination increased in areas that become more morphometrically dissimilar, or less hub-like with Δ*k <* 0, during adolescence. Conversely, macro-structural MRI features were positively correlated with adolescent change in MSN degree, i.e., cortical thickness, volume and surface area all decreased in regions that became less hub-like during adolescence. Here, we highlight regions that significant (*P*_*FDR*_ *<* 0.05) Δ*k* with grey outlines. (E) We estimated the correlation between the age effect on morphometric similarity and several prior maps of brain organization. We found a negative correlation between the effects of age on MSN nodal degree and several brain maps of metabolic rates, meaning that regions that showed decreases in degree of morphometric similarity tended to have increased metabolic rates. Conversely, the positive correlation between the age effect on MSN nodal degree and a map of cerebral blood volume means that regions that had decreased morphometric similarity over the course of adolescence had lower cerebral blood volume.

We did not find evidence for widespread sex differences in morphometric similarity during adolescence (**SI Fig. S8**), and while some regions showed significant effects of scanning site, these regions did not overlap extensively with regions that showed significant developmental changes in morphometric similarity (**SI Fig. S9**), the effects of site and age were not correlated (*ρ* = −0.07, *P* = 0.16). To address the potential impact of site-specific factors on internal replicability of our results, we split the sample by site into three internal replication datasets (347 scans acquired at WBIC, 98 at UCL and 33 at the MRC CBU). Despite the differences in site-specific sample size, we found a high spatial correspondence between our principal results on the whole NSPN cohort and the site-specific results derived from each of the 3 sub-cohorts (**SI Fig. S10**). 33 regional MSN nodes, primarily located in transmodal areas of the paralimbic (N=18) and heteromodal (N=10) cortex, had significant changes in weighted degree after correction for multiple comparisons (*P*_*FDR*_ *<* 0.05; see **SI Table 5, Fig. 3A, top**). We further confirmed that the directionality of significant age effects on morphometric similarity degree (*k*) was robust: (i) to sample size, using an ablation analysis which left out iteratively larger fractions of the data; and (ii) to composition of the sample, using a permutation approach that re-sampled subjects with replacement (**SI Fig. S11**). Lastly, we wanted to confirm that these between-subject, age-related changes in MSN degree were related to the developmental process of within-subject change in degree of similarity. However, we could not estimate within-subject linear rate of change as a random effect, because this would require at least 3 repeated measures per participant. Instead we simply estimated the difference in MSN degree at baseline and one-year follow-up at each cortical area for each of N=172 participants and found that this measure of within-subject longitudinal change was significantly positively correlated with between-subject age-related changes in MSN degree (Δ*k*; *ρ* = 0.41, *P*_*spin*_ *<* 0.001; see **SI Fig. 12**).

To test our hypothesis that development of morphometric similarity should be conditioned by the cytoarchitectonic features of cortex, we estimated the mean effect of age on weighted degree of all regions within each of four known cytoarchitectonic zones of cortex (Mesulam 2000; Mesulam 1998) (**Fig. 3B**). We found that iso-cortical or neocortical zones (idiotypic, unimodal and heteromodal) had decreased MSN degree, i.e. Δ*k <* 0, whereas paralimbic cortical areas had increased degree, i.e. Δ*k >* 0 (**Fig. 3B, SI Table S6**; for a breakdown of results by pre-defined functional sub-networks (Yeo et al. 2011) and further cytoarchitectonic classes (Economo and Koskinas 1925) refer to **SI Fig. S13**). We further observed overall decreasing within-zone similarity, whereas between-zone similarity was specifically and significantly increased between paralimbic and idiotypic cortex, as well as between paralimbic and heteromodal cortex (**Fig. 3C**). Moreover, we estimated changes in the connectivity between each individual cytoarchitectonic zone and the rest of the brain (**SI Fig. S15A**) and observed zone-specific patterns of re-organization in connectivity to the rest of the brain (**SI Fig. S11B**).

Lastly, we endeavoured to replicate our main findings from the NSPN cohort in an independent cohort. There was no other MRI dataset available to us that was directly comparable to NSPN in terms of providing repeated measures on MT. We instead used the Human Connectome Project Development (HCP-D) cohort (Somerville et al. 2018) as an approximate replication sample, which provides measures of the T1w/T2w ratio as an alternative proxy of intra-cortical myelination (Glasser and Van Essen 2011). We found consistent age-related decreases in cortical thickness, surface area and volume, and age-related increases in MT or T1w/T2w, across both HCP-D and NSPN cohorts. Most importantly, we found consistent results between cohorts in terms of age-related changes in degree of morphometric similarity, Δ*k*, across different cytoarchitectonic classes of cortex (*ρ* = 0.5, *P*_*spin*_ *<* 0.001,**SI Fig. 16, 17**). In MSNs derived from HCP-D data, as in MSNs from NSPN data, cortical areas in the paralimbic zone become increasingly similar, whereas areas of heteromodal association cortex become increasingly dissimilar, over the course of adolescence (see **SI Text** for details).

In an effort to understand the contribution of each of the six individual morphometric features to the adolescent change in morphometric similarity, we correlated the effect sizes of age (regional *t*-statistics) on the individual MRI phenotypes (**Fig. 2B**) with the age effects (regional *t*-statistic) on MSN weighted degree (**Fig. 3A**). We observed a divergent pattern: age-related changes in micro-structural micro-structural MRI markers were negatively correlated with adolescent changes in weighted degree (MD: *r* = −0.4, *P*_*spin*_ *<* 0.05; MT: *r* = −0.15, *P*_*spin*_ *<* 0.05; FA: *r* = −0.1), whereas macro-structural changes were positively correlated with adolescent changes in weighted degree (GM: *r* = 0.32, *P*_*spin*_ *<* 0.05; CT: *r* = 0.31, *P*_*spin*_ *<* 0.05; SA: *r* = 0.26, *P*_*spin*_ *<* 0.05) (**Fig. 3D**; **SI Fig. S18**). This result indicates that regions of unimodal, heteromodal and idiotypic cortex which became less morphometrically similar (or more morphometrically differentiated from the rest of the brain), as indexed by decreasing MSN degree during adolescence, tended also to become thinner and smaller, and more strongly myelinated (**Fig. 3D**). Thus the well-known adolescent processes of cortical thinning and increased myelination appeared to drive increasing morphometric dissimilarity or differentiation of neocortical nodes. Regions of paralimbic cortex that had increased MSN degree over the course of adolescence showed a similar pattern of decreased macro-structural and increased micro-structural metrics, but of smaller magnitude compared to isocortical areas, as indicated by *t*-values closer to zero (**Fig. 3D**).

### Biological and psychological context of adolescent changes in anatomical connectomes

We were interested in contextualizing age-related changes in MSNs in relation to prior maps of transcriptional and functional gradients, evolutionary change, and metabolic requirements (Sydnor et al. 2021). We found that the whole brain map of adolescent change in weighted degree of each node was significantly negatively correlated with commensurate maps of aerobic glycosis (*r* = −0.32; *P*_*spin*_ *<* 0.05) and the rates of oxygen (*r* = −0.44; *P*_*spin*_ *<* 0.001) and glucose metabolism (*r* = −0.48; *P*_*spin*_ *<* 0.001). Thus association and other isocortical nodes that had decreased MSN degree during adolescence tended to have increased metabolic demands in adulthood (**Fig. 3E**). Conversely, we found a positive correlation with a map of cerebral blood volume (*r* = 0.19; *P*_*spin*_ *<* 0.05), meaning that paralimbic regions that had increased MSN degree during adolescence tended to have decreased cerebral blood volume (**Fig. 3E**).

We also explored the psychological relevance of age-related changes in morphometric similarity. We conducted automated meta-analytic referencing using the NeuroSynth database of task-related fMRI activation co-ordinates (Yarkoni et al. 2011). This analysis revealed that isocortical regions that showed decreases in MSN degree (*t <* 0) were typically activated by tasks related to visual processing and imagery, motor control, and working memory. Conversely, paralimbic regions that showed increases (*t >* 0) in MSN degree were associated with self-evaluation of emotional content, nociception, and pain (**SI Fig. S14**).

### Adolescent development of structure-function coupling

We hypothesized that adolescent changes in brain structure, measured as increases or decreases of morphometric similarity, might change the strength of coupling between structural and functional brain networks (**Fig. 4A**). To test this hypothesis, we first estimated global structure-function coupling as the correlation between the ranked elements of the functional connectivity matrix and the morphometric similarity matrix for each subject, at each time-point (**Fig. 4B**). We modeled the linear effect of age on structure-function coupling using the same LME model as previously used for global, local and MSN metrics. We found that global structure-function coupling decreased over the course of adolescence (*t* = −5.04, *P <* 0.001; **Fig. 4C**), indicating a decoupling of functional connectivity from morphometric similarity.

**Fig. 4:**
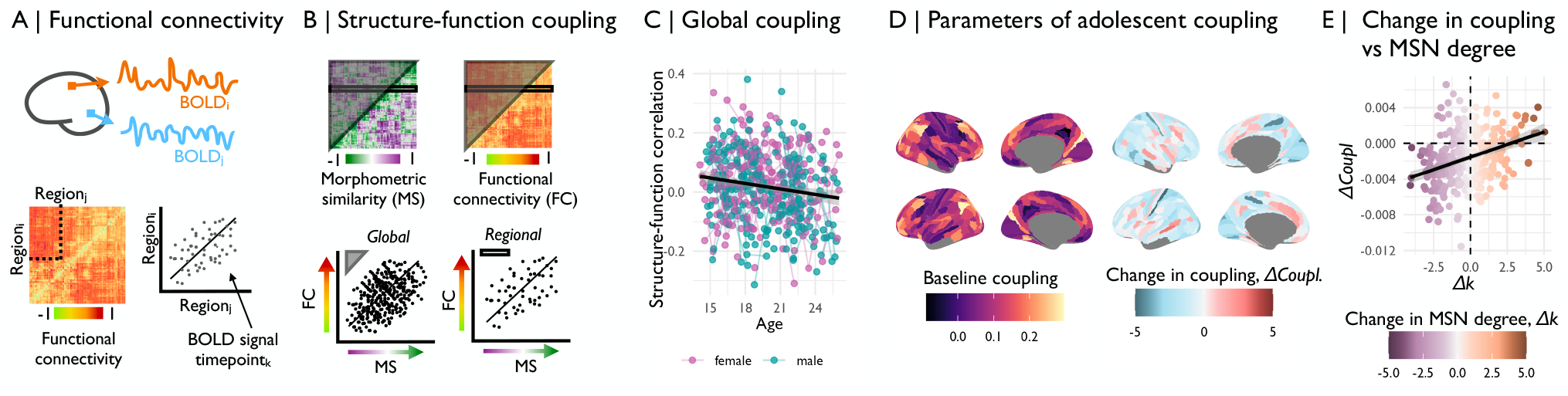
Adolescent development of structure-function coupling: (A) We derived a functional connectivity (FC) matrix, or functional connectome, for each scan by estimating the pairwise correlations between resting state fMRI time series averaged over all voxels in each of all possible pairs of two regions defined by the parcellation template. (B) We estimated global structure-function coupling by correlating the ranked edgewise structural and functional connectivity vectors derived from each participant’s FC matrix and MSN, respectively; and regional structure-function coupling as the correlation between the ranked vector of a region’s edges derived from the FC matrix and the MSN, respectively. (C) We estimated the linear effect of age on global structure-function coupling and found that there was a significant decline in coupling over the course of adolescence (*t* = −5.04, *P <* 0.001). We next estimated the linear effect of age on regional structure-function coupling using linear mixed effects models. From this analysis, we derived (D) a map of baseline structure-function coupling as the predicted coupling at age 14, and (E) a map of the rate of change in coupling, or the *t*-value of the effect of age on structure-function coupling. We found that 10 regions showed significant changes in structure-function coupling during adolescence, after correction for multiple comparisons (*P*_*FDR*_ *<* 0.05; **SI Fig. S12A**). (E) We found that the age effect on morphometric similarity was significantly positively correlated with the rate of change in coupling (*r* = 0.36, *P*_*spin*_ *<* 0.01). Thus, isocortical regions that become more morphometrically dissimilar (structurally differentiated) tended to have decreased strength of structure-function coupling over the course of adolescence.

We also tested this hypothesis regionally, estimating the correlations between the ranked elements of each row of the functional connectivity and morphometric similarity matrices (**Fig. 4B**), and then using the same LME to estimate age effects on regional structure-function coupling. From this analysis, we derived a map of baseline coupling, or the predicted coupling at age 14 years (**Fig. 4D, left**), as well as a map of adolescent changes in regional coupling (**Fig. 4D, right**). Baseline coupling was high in most iso-cortical areas, but lower in paralimbic areas (**Fig. 4D, SI Fig S19A, SI Table S7**). Coupling decreased most strongly in iso-cortical regions, and decreased less or increased slightly in paralimbic cortical regions (**Fig. 4E, SI Fig. S19B, SI Table S7**). It is notable that the majority of regions decreased in coupling over the course of adolescence (blue in **Fig. 4D**). We also found that structure-function coupling at baseline and the rate of change in coupling were negatively correlated (*r* = −0.35; *P*_*spin*_ *<* .001; **SI Fig. S20C**), thus regions that were more strongly coupled at baseline tended to have greater decreases in coupling over the course of adolescence.

We further investigated how baseline regional structure-function decoupling and its adolescent changes were related to baseline and adolescent changes in morphometric similarity. We found that baseline MSN degree at age 14 years (**SI Fig. S21**) was weakly correlated with baseline structure-function coupling (*r* = 0.15, *P*_*spin*_ *<* 0.05; **SI Fig. S20B**), such that regional hubs at baseline had stronger structure-function coupling. There was also significant positive correlation between adolescent changes in MSN degree and structure-function coupling (*r* = 0.36, *P*_*spin*_ *<* 0.01; **Fig. 4F**), meaning that isocortical regions which became more dis-similar from the rest of the brain during adolescence had weaker structure-function coupling, whereas paralimbic regions which became more morphometrically similar had stronger coupling (**Fig. 4F, SI Fig S20**).

### Co-development of structural and functional network changes

In order to further understand how adolescent changes in structural brain networks relate to concomitant changes in brain functional networks, we estimated age-related changes in multiple network metrics (**Fig. 5A**). We did not find significant associations between adolescent changes in morphometric similarity and adolescent changes in weighted degree of functional connectivity (*r* = −0.05, *P*_*spin*_ = 0.4; **SI Fig. S22, Fig. 5A**) or multiple other network metrics, including within- and between-network connectivity, eigenvector centrality, clustering coefficient, and efficiency (**Fig. 5A**). However, motivated by prior work (Baum et al. 2020), we were particularly interested to assess whether developmental changes in morphometric dissimilarity were associated with changes in participation coefficient, a measure of the topological diversity of functional connectivity between functionally specialized modules (Yeo et al. 2011; Meunier et al. 2010) (**Fig. 5B**). Regions with a high participation coefficient have a relatively high proportion of inter-modular connections to nodes in other modules, thus they may have the capacity to integrate information across multiple sub-graphs or modules of the whole brain connectome and have been designated as “connector hubs”. Conversely, regions with a low participation coefficient have more locally segregated connectivity within their respective modules and have previously been designated “provincial hubs” because of their important role in communication between functionally specialised modules (Fornito et al. 2016). We found that adolescent increases in regional participation coefficient were largely located in isocortical regions and decreases were more concentrated in paralimbic regions, as well as medial prefrontal regions (**Fig. 5C**). Further, we found that adolescent changes in MSN degree were correlated with adolescent changes in PC (*r* = −0.24, *P*_*spin*_ *<* 0.01, **Fig. 5A**, *bottom*), such that isocortical regions that became more morphometrically dissimilar over the course of adolescence had increased PC over the same period (**Fig. 5D, SI Fig. S23, SI Table S8**), whereas paralimbic regions that became more morphometrically similar had decreased PC during adolescence. We thus established that increases in morphometric dissimilarity, or structural differentiation from the rest of the brain, were associated with increasing diversity of functional connectivity, measured as a relative strengthening of inter-modular connectivity, potentially representing an increased ability to integrate information across multiple, structurally differentiated and functionally specialised modules.

**Fig. 5:**
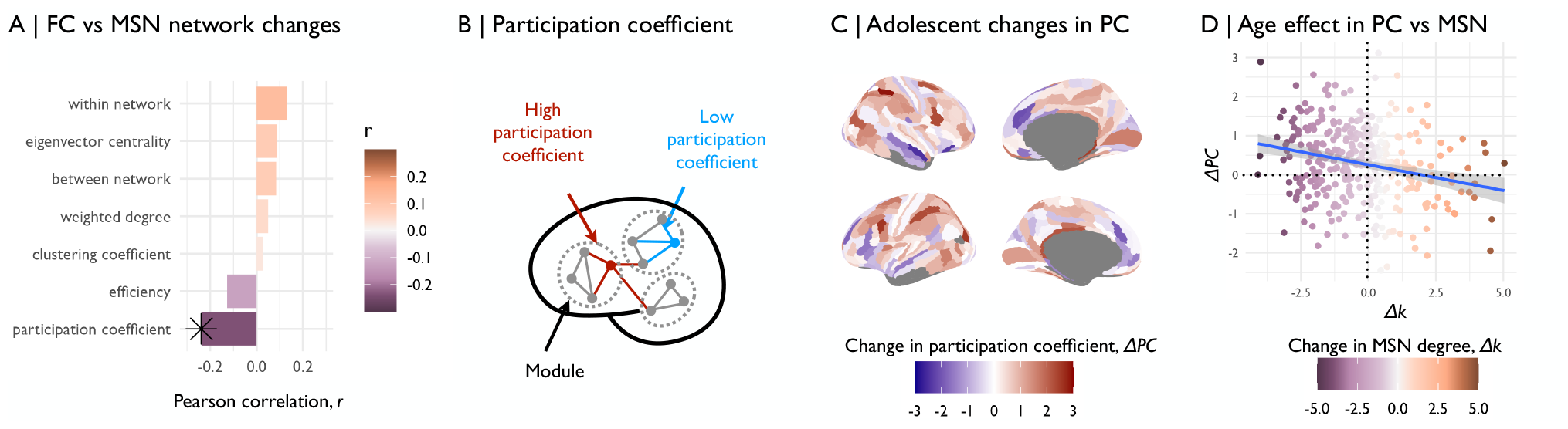
Morphometric dissimilarity was associated with functional participation: (A) We estimated age-related changes in multiple network metrics of regional nodes in fMRI connectomes over the course of adolescence. We then correlated the adolescence-related changes in these fMRI network phenotypes with changes in morphometric similarity. We found that only the participation coefficient (PC), a measure of inter-modular connectivity, showed adolescent changes that were significantly associated with adolescent changes in MSN degree. (B) We then estimated the regional participation coefficient (PC) at each node for each connectome. PC is measured as the ratio between a node’s intra-modular degree (edges connecting to other nodes in the same functional module (Yeo et al. 2011; Meunier et al. 2010)) and its inter-modular degree (edges connecting to nodes in other modules). (C) We estimated the linear effect of age on PC. Functional participation increased over the course of adolescence in association cortical regions and decreased in primary motor and sensory regions, as well as medial prefrontal regions. (D) We estimated Spearman’s correlation between regional age-related changes in morphometric similarity and regional age-related changes in functional participation coefficient. We found that regions that became more morphometrically dissimilar over the course of adolescence tended to increase in their functional participation.

## Discussion

We set out to investigate the hypothesis that there are developmental changes in human brain structural networks during adolescence, and that cortical areas may become more or less hub-like in the connectome depending on their maturing cytoarchitectonic differentiation from the rest of the cortex. We used morphometric similarity analysis of multiple MRI metrics to measure brain anatomical networks from repeated structural MRI assessments of a large cohort of healthy young people. We discovered that there were indeed significant developmental changes in the hubness of cortical areas during adolescence, and that the regional trajectories of adolescent change in MSN hubness or degree, Δ*k*, were remarkably distinct between paralimbic and all other (isocortical) zones of cortex.

According to the cytoarchitectonic scheme defined by Mesulam (Mesulam 1998; Mesulam 2000), isocortex or neocortex is defined by 6 cortical layers, it encompasses the majority of human cortex, and it can be sub-divided into three zones of idiotypic, unimodal and heteromodal cortex. We found that almost all such isocortical areas became less similar or more dissimilar from the rest of the cortex during adolescence, and therefore less hub-like in morphometric similarity networks. The most likely interpretation of this process of “de-hubification” is that each of these areas is becoming more structurally differentiated from the rest of the brain, more unique in its cytoarchitectonic or myleoarchitectonic organization. We found that isocortical regions with decreasing hubness typically had more rapidly shrinking cortical thickness, volume and surface area, and greater increases in myelination indexed by MT. Coupled changes in cortical shrinkage and cortical myelination have been well-replicated in prior neurodevelopmental MRI studies of childhood and adolescence (Whitaker et al. 2016; Mills, Goddings, Herting, et al. 2016) and can be interpreted as a process of pruning connections at a cellular level, and consolidating those connections that survive by myelination. It therefore seems likely that the increasing dis-similarity of isocortical nodes in morphometric similarity networks reflects their increasingly selected and distinctive profile of (myelinated) anatomical connections to the rest of the brain.

The paralimbic zone is defined by having less than 6 cortical layers, and lacking a well-defined layer 4 of granule cells. It represents a minority of total cortex, including insular, orbito-frontal, temporal polar, and cingulate cortices. Cytoarchitectonically and connectionally, the paralimbic zone is regarded as transitional or intermediate between the most phylogenetically primitive, 3-layered regions of allocortex, e.g., hippocampus or pyriform cortex, and the 6-layered neocortex or isocortex. We found that almost all paralimbic areas became more morphometrically similar, or more hub-like, over the course of adolescence. There are at least two plausible interpretations of this result: local (absolute) or contextual (relative) change in cortical structure. Locally, it could be that paralimbic cortex becomes “less primitive” or cytoarchitectonically more similar compared to isocortex during adolescence. Across the insula of the primate brain, for example, there is an antero-posterior gradient of cortical architectonics from the agranular, peri-allocortical organization of anterior (rostral) insula, continuous with 3-layered pyriform cortex, to the dysgranular, pro-isocortical organization of posterior (caudal) insula, continuous with 6-layered heteromodal and unimodal association areas of temporal and parietal cortex (Mesulam and Mufson 1982). This cytoarchitectonic gradient has evolved phylogenetically from reptiles to primates, and it is conceivable that this process might be recapitulated ontogenetically, with an increasing proportion of the insula becoming organized more like isocortex and less like allocortex over the course of development. This would be consistent with our observations of greater morphometric similarity between isocortical (idiotypic and heteromodal) and paralimbic zones, and therefore higher degree of paralimbic cortical nodes, over the course of adolescent MSN development. An alternative interpretation is that the paralimbic cortex becomes relatively more similar to isocortex, as the isocortical areas become relatively more dissimilar to (differentiated from) each other. In other words there may be no local cytoarchitectonic maturation of progressively more isocortical lamination but paralimbic areas could nonetheless appear to become more similar “on average” across the brain as the corollary of isocortical or neocortical differentiation associated with adolescent shrinkage and myelination. Further studies linking MRI metrics and morphometric similarity to underlying cellular changes in development of mammalian cortex will be required for definitive mechanistic resolution of the increasing hubness of paralimbic cortex, as well as to confirm the pruning-and-myelination mechanism proposed to account for decreasing hubness of isocortex during adolescence. Indeed, prior work in children has linked morphometric dissimilarity between isocortical and paralimbic zones to better cognitive performance, suggesting that structural differentiation is a prerequisite for healthy development in the pre-adolescent period, a finding which supports the relevance of understanding changes in morphometric similarity during development (Wu et al. 2022).

We proceeded to investigate the consequences for brain functional connectivity of this cytoarchitectonically-aligned divergence in structural network development, and made two interesting observations. First, we found that morphometric similarity and functional connectivity, measured in the same scanning session, were significantly but modestly coupled at baseline (*r* ≤ 0.25 at 14 years) and the strength of structure-function coupling globally declined over the course of adolescence, as previously reported (Zamani Esfahlani et al. 2022). However, at a regional level, there was again some evidence for cytoarchitectonically aligned divergence in developmental trajectories. Isocortical areas tended to have reduced structure-function coupling over the course of adolescence compared to paralimbic areas. And adolescent change in structure-function coupling at each region was positively correlated with the adolescent change in MSN degree that was previously shown to be cytoarchitectonically aligned. Second, we found that the participation coefficients for each regional node in the fMRI connectomes were also developmentally divergent on cytoarchitectonic lines: most isocortical areas had increased participation and most paralimbic areas had decreased participation; and adolescent change in participation was positively correlated with the cytoarchitectonically aligned adolescent change in MSN degree. Collectively these results suggest that as isocortical areas become less hub-like or more structurally differentiated their pattern of functional interactions with other cortical areas becomes less constrained or more diverse. Morphometric similarity is a proxy for (often reciprocal) monosynaptic connectivity between areas of same cytroarchitectonic class, whereas functional connectivity is often regarded as a proxy for polysynaptic connectivity. Low strength of structure-function coupling and high participation means that mature isocortical areas can interact functionally with other areas even if they are cytoarchitectonically dis-similar or affiliated to different modules of the fMRI connectome, possibly relying more on polysynaptic (indirect) axonal connections or circuit-level modulation of neuronal activity (Baum et al. 2020). Given prior theories on the importance of network integration and segregation to different aspects of cognition (Baum et al. 2020), this developmental shift of heteromodal and other isocortical nodes to increasing diversity of functional connectivity and increasing differentiation of cortical structure could be relevant to the emergence of a wide range of domain-general or “higher order” cognitive skills which depend on integrative network properties (Crossley et al. 2013). The opposite trend demonstrated by some (mostly paralimbic) areas - towards increased structure-function coupling and decreased participation - convergently suggests consolidation of intra-modular interactions between morphometrically similar areas specialised for a specific function, e.g., interoception, that emerges during adolescence. Further work will be needed to test these predictions of the cognitive consequences of cytoarchitectonically aligned changes in MSN and fMRI network connectivity.

There are some notable methodological issues. First of all, the test-retest reliability of MSNs has not previously been assessed. Here, we estimated the longitudinal between-person rank stability as a proxy and found reasonable reliability, with baseline and follow up regional between-person ranks positively correlated in 241 regions (see **Supplementary Text** and **SI Fig. S24**). Future work will have to assess test-retest reliability more directly. Only a minority of cortical areas demonstrated significantly non-zero changes in similarity during adolescence yet most of our analysis has included data from all cortical areas, e.g., to investigate differences in development of similarity between cytoarchitectonic classes or cortical types. In doing so we assumed that developmental change in similarity was widespread throughout the brain but our statistical power to demonstrate significant change at a regional level was constrained by the number of participants, the number of scans per participant, the nominal degrees of freedom for estimation of similarity in each scan, and the multiple comparisons correction needed to control *FDR <* 5% for all 358 cortical regions tested. We expect that future studies of brain similarity development will achieve greater statistical power by addressing some or all of these constraints. Further, to date there is no consensus on how best to estimate the coupling between structural and functional connectivity, and various methods have been used to define both structural and functional networks (Baum et al. 2020; Zamani Esfahlani et al. 2022; Liu et al. 2022), which may contribute to the lack of consistency in the overall pattern of previously reported results. Notably, structure has so far been defined from DWI networks (Baum et al. 2020) or graph theoretical properties of such networks (Zamani Esfahlani et al. 2022). However, it is conceivable that new insights can be gained into how structure constrains function by employing different structural network modeling approaches, including more directly modelling relevant maturational processes of increasing myelination paired with cortical thinning, as is possible using morphometric similarity networks.

Overall, we conclude that adolescence is associated with an extensive developmental programme of increasing structural differentiation and functional integration of isocortical zones, and increasing structural similarity and functional segregation of paralimbic cortex .

## Methods

This study included data from an accelerated longitudinal study (Kiddle et al. 2017) of adolescents ages 14 to 26 years (51 % female) who were invited to undergo functional and structural neuroimaging assessments on at least two accasions: at baseline and at a one year followup assessment, with a subset of the sample invited to come in six months after baseline for an additional scan (**SI Fig. S1**). Participants provided informed written consent for each aspect of the study, and parental consent was obtained for those aged 14–15 years. .The study was ethically approved by the National Research Ethics Service and conducted in accordance with U.K. National Health Service research governance standards (see **SI Appendix Text** for further details).

### Structural MRI acquisition and preprocessing

The MRI data were acquired using a multi-parametric mapping (MPM) sequence (Weiskopf et al. 2013) at three sites, on three identical 3T Siemens MRI scanners (Magnetom TIM Trio, VB17 software version) with a standard 32-channel radio-frequency (RF) receive head coil and RF body coil for transmission. The anatomical, diffusion weighted and functional imaging data were collected during the same session. The anatomical MRI data were acquired using a single-shot echo planar imaging sequence (63 gradient directions with b-value = 1000 mm/s^2^ and 5 unweighted B0 images) was used to acquire a HARDI with the following scanning parameters: slice number = 70 consecutive; slice thickness = 2 mm; field of view = 192 × 192 mm; TE = 90 ms; TR = 8700 ms; voxel size = 2.0 mm isotropic.

We pre-processed the anatomical data using the recon-all command in Freesurfer v5.3.0 (Fischl 2012). In short, the pipeline included the following steps: non-uniformity correction, projection to Talairach space, intensity normalisation, skull stripping, automatic tissue and subcortical segmentation, and construction of smooth representations of the gray/white interface and the pial surface. Subsequently, the DWI volumes were aligned to the R1 image for each subject. Lastly, we parcellated the anatomical and DWI scans into 360 bilateral parcels, using the Humman Connectome Project (HCP) parcellation atlas (Glasser, Coalson, et al. 2016).

### fMRI acquisition and pre-processing

The functional MRI data were acquired using a multi-echo (ME) echo-planar imaging sequence with the following scanning parameters: TR = 2.42 s; GRAPPA with acceleration factor = 2; flip angle = 90°; matrix size = 64 × 64 × 34; field of view = 240 mm by 240 mm; inplane resolution = 3.75 mm by 3.75 mm; slice thickness = 3.75 mm with 10% gap, with sequential acquisition of 34 oblique slices; bandwidth = 2368 Hz/pixel; and echo times (TE) = 13, 30.55, and 48.1 ms.

The pre-processing pipeline has been described in depth in prior publications (Váša, Romero-Garcia, et al. 2020; Dorfschmidt et al. 2022). Briefly, this included: multi-echo independent component analysis (ME-ICA) to remove non-BOLD components (Kundu, Inati, et al. 2012; Kundu, Brenowitz, et al. 2013); Cerebrospinal fluid (CSF) regression using Analysis of Functional NeuroImages software (AFNI; (Cox 1996)); parcellation into 360 bilateral cortical regions using the HCP template (Glasser, Coalson, et al. 2016); band-pass filtering (frequency range 0.025 to 0.111 Hz); removal of 30 dropout regions, defined by a low *Z* score of mean signal intensity in at least one participant (*Z <* −1.96); functional connectivity esusing Pearson’s correlation between all possible combinations of regional timeseries; and Fisher’s *r* − *Z* transformation. Finally, to remove any residual effects of head motion on functional connectivity, we regressed each pairwise correlation between regions on the time-averaged head motion of each participant (mean framewise displacement). We retained the residuals of this regression, i.e., motion-corrected *Z* scores, as the estimates of functional connectivity for this analysis.

### Morphometric feature estimation and quality control

We derived FreeSurfer’s standard morphometric features: CT, GM, SA. Previous work on this sample had indicated that MT adolescent changes with age were most pronounced at 70% cortical depth from the pial surface (Whitaker et al. 2016); thus regional MT values were estimated at that depth. Lastly, regional volumes for fractional anisotropy (FA) and mean diffusivity (MD) were derived from the DWI scans.

We excluded 11 subjects due to outliers in at least one morphometric feature (*MAD* ≥ 5; see **SI Fig. S2** and **SI Appendix Text** for further details). 2 regions were excluded due a local signal dropout, defined as *MAD* = 0 in at least one morphometric feature across subjects, which led to the exclusion of two regions (*L_H, R_H*), such that the total number of regions analysed henceforth was 358 (see **SI Fig. S3** and **SI Appendix Text** for further details).

### Modeling of developmental change in morphometric features

We estimated linear age-related changes or development in six morphometric features at global scale, i.e., on average for each feature over all regions, and locally, for each feature *f* at each region (*i* = 1…*N* = 358), using linear mixed effects models, with a fixed effect of age, sex and site, and a random effect for the repeated measures on each participant, as follows:

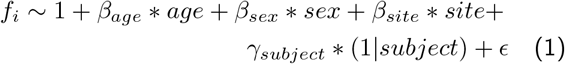

where *f*_*i*_ refers to the morphometric feature value at region *i, β* refers to the coefficients for the fixed effects, *γ*_*subject*_ refers to the coefficients for the random effect, and *ϵ* represents the residual error.

### Adolescent changes in morphometric similarity

We derived subject-specific structural connectomes, i.e. morphometric similarity networks. To this end, we standardized each morphometric feature within each subject using MAD (Váša, Hobday, et al. 2022). We then estimated morphometric similarity networks for each subject by calculating the Pearson correlation between their standardized feature vectors for each possible pair of regions. This resulted in a 358*×*358 symmetric matrix, indicative morphometric similarity between cortical regions.

We first estimated regional morphometric similarity, or weighted degree *k* as the mean across a region’s edges. Then we estimated the linear effect of age on MSN weighted degree, using linear mixed effects models (**Fig. 1G**) with a fixed effect of age, sex, and site and random effect of subject, as follows:

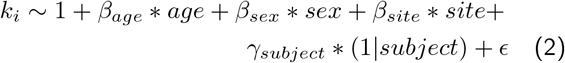

where *k*_*i*_ refers to the morphometric similarity strength, or weighted degree, of regional node *i, β* refers to coefficients for the fixed effects, *γ*_*subject*_ refers to the coefficients for random effects, and *ϵ* represents the residual error. From this model, we estimated the adolescent rate of change in morphometric similarity, or the age effect on weighted degree at each node of MSN, as the *t*-statistic of the age effect.

We tested for the robustness of the derived age-related changes in morphometric similarity using two sensitivity analyses: First, we estimated the correlation between age effects derived using an ablation analysis of ever-decreasing sample sizes, i.e. we randomly sampled decreasing numbers of subjects and re-estimated the effect of age on the smaller samples. Secondly, we derived confidence intervals around the estimated age effects in a leave-N-out bootstrap analysis, where we randomly left out 10% of the sample in 1000 permutations and re-estimated the age effects on morphometric similarity (see **Supplementary Fig. 8** for details).

In order to decode the regional changes in morphometric similarity by cell type and functional modules, we averaged weighted degree over all regions within each cytoarchitectonic zone and functional network of cortical areas defined *a* priori by the respective reference brain atlas (Economo and Koskinas 1925; Mesulam 1998; Mesulam 2000; Yeo et al. 2011).

We then estimated the correlation between age-related changes (*t*-values) in individual morphometric features, estimated by **Equation 1**, and age-related changes (*t*-values) in morphometric similarity, estimated by **Equation 2**. Each analysis of spatial co-location or correlation between two cortical maps was reported with both the parametric *P*-value corresponding to the Pearson correlation (*r*), as well as a *P*-value derived from the more conservative “spin-test” permutation. Spatial autocorrelation of statistical brain maps can cause inflated estimates of the probability of spatial co-location or correlation between two maps (Alexander-Bloch et al. 2013; Váša, Seidlitz, et al. 2018). The spin test procedure addresses this issue by conserving the spatial autocorrelation of the maps by randomly “spinning” or spherically rotating each map 1000 times over the surface of the brain and calculating the spatial co-location statistic after each spin permutation.

We estimated the anatomical co-location of the map of age-related changes in morphometric similarity with various maps of cortical organization, including metabolic rates, blood volume, and functional hierarchy (Sydnor et al. 2021). To do this, we correlated the ranked map of age-related changes in morphometric similarity with each prior map, and then estimated the significance of the correlation while controlling for spatial auto-correlation using a spin-test (Váša, Seidlitz, et al. 2018).

We further assessed the psychological relevance of the map of age-related changes in morphometric similarity using Neurosynth, an automated meta-analytical tool (Yarkoni et al. 2011). We generated a volumetric version of the regional map of adolescent changes in morphometric similarity (code available at https://github.com/LenaDorfschmidt/neurosynth_analysis) and uploaded it for automated comparison to the Neurosynth database (https://neurosynth.org) of task-related fMRI activation coordinates, which returned the correlation values of the map with a wide set of terms related to fMRI task activation experiments.

### Adolescent changes in structure-function coupling

We estimated global structure-function coupling as the Spearman correlation between the upper triangle of each subjects’s structural (MSN) and functional connectivity (FC) networks at each timepoint (**Fig. 4A,B**). Local structure-function coupling was estimated at each node as the Spearman correlation between the node’s edges in the structural and functional networks (**Fig. 4A,B**).

Then, we estimated parameters of adolescent change in structure-function coupling. Specifically, we estimated the linear effect of age on regional structure-function coupling strength using linear mixed effects models, with a fixed effect of age, sex and site, and a random effect of subject, as above for MSN regional strength (**Equation 2**). From this model, we proceeded to derive the local structure-function coupling at baseline (age 14) and the rate of change in coupling over the course of adolescence, as the *t* − *values* of the effect of age (see **SI Appendix Text** for further details).

### Co-location with adolescent changes in functional diversity

We assessed whether changes in morphometric similarity during adolescence were associated with changes in functional brain networks which might represent adolescent changes in cognition and behavior. We estimated multiple network metrics on the unthresholded functional connectomes (**Fig. 5A**) and used linear mixed effects models as above to estimate the linear effect of age on each network metric (see **SI Appendix Text** for further details).

## Data and Code

The raw data is publicly available at https://nspn.org.uk and the processed data will be made available upon publication. The code can be accessed at https://github.com/LenaDorfschmidt/NSPN-MSN.git.

