## Supplementary material for "Human adolescent brain similarity development is different for paralimbic versus neocortical zones": SI

### Contents

**Data and pre-processing** **4**
Participants . . . . . 4
Morphometric feature estimation and quality control . . . . . 4

**Adolescent changes in morphometric similarity** **4**

**Biological and psychological context of adolescent changes in anatomical connectomes** **5**

**Adolescent changes in structure-function coupling** **5**

**Co-location with adolescent changes in functional diversity** **5**

**Internal replication of results in site-specific NSPN sub-cohorts** **6**

**Replication of results for in an independent dataset** **6**

**Rank-stability of morphometric similarity networks** **7**

**NSPN Consortium Author List** **8**

### Data and pre-processing

#### Participants

The data used in this work was collected as part of the Neuroscience in Psychiatry Network (NSPN), a joint initiative by the University of Cambridge and University College London (Kiddle et al. 2017), with the aim of using an accelerated longitudinal design to measure developmental brain changes in a sample drawn from the population of Greater London and Cambridgeshire that was broadly representative of the populations of England and Wales (**SI Fig. 1**). A total of 306 adolescents aged 14 to 26 years (51 % female) were invited to undergo functional and structural neuroimaging assessments. The exclusion criteria for this subsample included a current or past history of neurological disorders, current treatment for psychiatric disorder or drug or alcohol dependence, as well as learning disabilities. Each participant was invited to provide data on at least two occasions; at baseline and at a one year follow-up assessment, with a subset of the sample invited to come in six months after baseline for an additional scan. The cohort was scanned a total of 556 times. The study was ethically approved by the National Research Ethics Service and conducted in accordance with U.K. National Health Service research governance standards.

#### Morphometric feature estimation and quality control

We derived FreeSurfer's standard morphometric features: CT, GM, SA, IC , MC, CI, FI. Previous work on this sample had indicated that MT adolescent changes with age were most pronounced at 70% cortical depth from the pial surface (Whitaker et al. 2016); thus regional MT values were estimated at that depth. Lastly, regional volumes for fractional anisotropy (FA) and mean diffusivity (MD) were derived from the DWI scans.

In order to identify potential outliers, first we standardized each (global) morphometric feature using the non-parametric metric median absolute deviation (MAD) (Leys et al. 2013). We normalized each features across nodes within each scan, i.e.:

$$MAD_{f,s} = \frac{X_{f,s} - median(X_{f,s})}{k * median(|X_{f,s} - median(X_{f,s})|)} \tag{1}$$

where $X_{f,s}$ is the vector of regional feature values for a single feature $f$ across regions for a single scan $s$, and $k \approx 1.4826$ is a constant, which ensures that for large $N$ the median absolute deviation is approximately equal to the standard deviation. Thus $MAD_{f,s}$ is a vector of standardized feature values for a single feature $f$ across regions for a single scan $s$.

We excluded 11 subjects due to outliers, with $MAD \geq 5$ set as the threshold based on visual interpretation of the distributions of data, in at least one global morphometric measure (**SI Fig. 2**).

We then estimated MAD locally, as in **Equation 1**, for each morphometric feature across subjects. First, we excluded regions with signal dropout, defined as $MAD = 0$, which led to the exclusion of two regions (*L_H, R_H*), such that the total number of regions analysed henceforth was 358. Next, within each subject, we excluded all regions with $MAD \geq 5$ (**SI Fig. 3**). At this step of the quality control pipeline, we found that the curvature features, MC, IC, CI, and FI, demonstrated much larger numbers of outliers across all regions (**SI Fig. S3**). Prior work has convincingly demonstrated that MSNs are stable to the use of varying feature sets Seidlitz et al. 2018; Fenchel et al. 2020, as well as to reductions in the number of features used for estimation of similarity (correlation) between regions Seidlitz et al. 2018. As expected, morphometric similarity is more precisely estimated on the basis of more degrees of freedom, i.e., more regional mean MRI metrics in the feature vector for each regional node. When MSNs were estimated on the basis of 10 features, and compared to the MSNs estimated on the basis of a sub-sample of 5 features, the two networks were highly correlated with each other but the 5-feature network had much higher between-subject variability Seidlitz et al. 2018. Therefore, we know that restricting feature selection will increase variability and reduce precision of similarity estimation. However, this is not the only consideration; indeed it is trumped by data quality considerations. Estimating correlations on the basis of 10 or fewer nominal degrees of freedom is clearly vulnerable to bias by one or more outlying variables, and quality control identified that several of the available MRI metrics had frequent outlying observations. We therefore excluded these "noisier" metrics of curvature and gyrification and focused on a subset of 5 metrics, accepting the cost in degrees of freedom was worth it in terms of data quality control.

### Adolescent changes in morphometric similarity

We estimated regional morphometric similarity, or weighted degree as the mean across a region's edges as follows:

$$s_i = \sum_{j=1;j\neq i}^{N} w_{i,j} \tag{2}$$

where $s_i$ is the mean weighted degree of node $i$, $N$ is the number of nodes in the network, and $w_{i,j}$ is the weight of the edge between node $i$ and an arbitrary node $j$. The sum is taken over all edges $w_{i,j}$ $(j \neq 1, 2, 3, \ldots N)$.

### Biological and psychological context of adolescent changes in anatomical connectomes

#### Co-location with prior fMRI task activation

We performed automated meta-analytic referencing of the thresholded map ($P_{FDR} < 0.05$) of adolescent changes in MSN degree, $\Delta k$, using the NeuroSynth database Yarkoni et al. 2011. To this end, we generated a volumetric version of the thresholded $\Delta k$ map by. Specifically, we assigned the $\Delta k$ value of each of the N=33 significant ($P_{FDR} < 0.05$) regions to its respective parcel in the volumetric nifti file of the parcellation. We uploaded this map to the online database. The NeuroSynth decoder registered our map with its set of cognitive terms and their coordinates in standard space generated through automatic parsing of the literature.

The decoder then returned a ranked list of cognitive terms with their correlation values describing the strength of association to the map of adolescent changes in morphometric similarity. The correlation values generated indicate whether a given term was positively or negatively associated with our map, i.e. a positive correlation between a NeuroSynth term and our map indicates that this term is more often mentioned in studies that show activation in regions where we observe a positive $\Delta k$. Conversely, a negative correlation value indicates that a given term is more frequently mentioned in studies that see activation in regions where we observe a negative $\Delta k$. It is worth mentioning that this automated process of generating meta-analytical maps is not perfect, i.e. each individual coordinate extraction may not be faultless, however, the number of articles parsed results in highly accurate meta-analytical maps.

#### Analysis of adolescent changes in MSN degree by Mesulam zones

We analysed changes in morphometric similarity by Mesulam zones Mesulam 1998; Mesulam 2000, distinct regions of the cerebral cortex characterized by variations in cellular composition and organization. Prior work has demonstrated that cytoarchitectonically similar regions are often involved in similar functions Mesulam 2000.

We focused on a cytoarchitectonic classification of similarity, and developmental changes in similarity, because the cytoarchitectonic and myeloarchitectonic similarities and differences between cortical areas seem most likely to be directly represented by the similarity or dissimilarity of their macro- and micro-structural MRI feature vectors. However, it is also conceivable there could be coordinated developmental changes of similarity between cortical areas constituting nodes of more broadly defined anatomical systems.

### Adolescent changes in structure-function coupling

We estimated parameters of adolescent change in structure-function coupling. Specifically, we estimated the linear effect of age on regional structure-function coupling strength using linear mixed effects models, with a fixed effect of age, sex and site, and a random effect of subject, as follows:.

$$CS_i \sim 1 + \beta_{age} * age + \beta_{sex} * sex + \beta_{site} * site + \gamma_{subject} * (1|subject) + \epsilon \quad (3)$$

where $CS_i$ refers to the strength of structure-function coupling at region $i$, $\beta$ refers to the coefficients for the fixed effects, $\gamma_{subject}$ refers to the coefficients for random effects, and $\epsilon$ represents the residual error.

We proceeded to derive the local structure-function coupling at baseline (age 14) as the predicted coupling value from **Equation 3**, i.e.,

$$CS_{14_i} = 1 + 14 * \beta_{age} + 0.5 * \beta_{sex} + 1/3 * \beta_{site}; \quad (4)$$

and the rate of change in coupling over the course of adolescence, as the $t - values$ of the effect of age, $\beta_{age}$, estimated by **Equation 3**.

### Co-location with adolescent changes in functional diversity

To this end we estimated multiple network metrics on the unthresholded functional connectomes (**Fig. 6A**), including the weighted degree, within- and between functional network connectivity using pre-defined functional networks (Yeo et al. 2011), the clustering coefficient, efficiency and participation coefficient (**Fig. 6B**), a measure of functional diversity estimated as the inter-modular connectivity mediated by each node. We then used linear mixed effects models to estimate the linear effect of age on the participation coefficient in each region $i$ as follows:

$$PC_i \sim 1 + \beta_{age} * age + \beta_{sex} * sex + \beta_{site} * site + \gamma_{subject} * (1|subject) + \epsilon \quad (5)$$

where $PC_i$ refers to the participation coefficient at region $i$, $\beta$ refers to the coefficients for the fixed effects, $\gamma_{subject}$ refers to the coefficients for random effects, and $\epsilon$ represents the residual error.

### Internal replication of results in site-specific NSPN sub-cohorts

The NSPN sample included scans from three sites: The NSPN sample included scans from three sites: Wolfson Brain Imaging Center Cambridge (WBIC, N = 347); University College London (UCL; N = 98); Cambridge MRC Cognition and Brain Sciences Unit, Cambridge (CBSU; N = 33). In our main analysis, we correct for the effect of scanning site using a fixed effect of site (see **SI Fig 9**). The multi-site nature of the dataset however also allows us to assess the sensitivity of our main results to scanning site. To this end, we split the NSPN by site and subsequently estimated adolescent changes in morphometric similarity in each site independently. We find that the resulting maps of adolescent changes in morphometric similarity (**SI Fig 10**) are significantly positively correlated with the original map using the full sample ($P_{spin} < 0.001$; $\rho_{WBIC} = 0.92; \rho_{CBSU} = 0.37, \rho_{UCL} = 0.63$).

### Replication of results for in an independent dataset

#### Data

We determined the replicability of our main results in an independent dataset. Our main sample, the NSPN dataset, is unique in various aspects: (i) it is an accelerated longitudinal design, offering the opportunity to study longitudinal effects over the entire course of adolescence; (ii) it is one of very few adolescent multi-modal datasets that includes DWI in this age span, and to the best of our knowledge the only public adolescent dataset that used included magnetization transfer images. The latter is of particular interest for this study given the known process of adolescent changes in myelination. We were unable to find an adolescent dataset that fulfilled both these criteria.

Nevertheless, we attempted a replication of our main results from the NSPN sample in the Human Connectome Project Development (HCP-D) sample Somerville et al. 2018, a cross-sectional cohort of children and adolescents aged 5-21 years (**SI Fig. 16A**), for which multimodal imaging data, including T1 and T2 images, as well as DWI data were acquired. For better comparison with the NSPN sample, we included only HCP-D subjects aged 14 and older (N=334).

#### Preprocessing of HCP-D data

DWI data were processed using QSIprep v.0.16.1 Cieslak et al. 2021, an automated preprocessing pipeline that handles distortion correction, motion correction, denoising, coregistering and resampling of DWI scans. We used a containerized version available at: https://hub.docker.com/r/pennbbl/qsiprep/. T1w and T2w images were processed using FreeSurfer v.6.0.1 within a containerized version of fMRIprep v.20.2.4 available at https://hub.docker.com/r/nipreps/fmriprep/ Esteban et al. 2019. Since the HCP-D dataset does not include MT images, we constructed T1w/T2w ratio from the raw T1- and T2-weighted images Glasser and Van Essen 2011. The T1w/T2w ratio has been shown to enhance the contrast to noise ratio for myelin and may thus function as a comparable measure to MT Glasser and Van Essen 2011. First, we used a rigid body registration of the T2w image to the T1w image using FSL's FLIRT Jenkinson et al. 2002 with 6 parameters and the mutual information cost function. Subsequently, we resampled the T2w image using the spline interpolation algorithm of FSL's applywarp tool to minimize the white matter and CSF contamination of gray matter voxels that would result from the volumetric blurring inherent in trilinear interpolation. We then divided the T1w image by the T2w at each voxel, and parcellated each image into 360 bilateral regions using the HCP parcellation Glasser, Coalson, et al. 2016.

We excluded 30 subjects as outliers in one or more global morphometric features as defined by a global feature value of $MAD > 5$. The final sample consisted of 304 subjects (151 females).

#### Comparison of HCP-D and NSPN results

First, we modelled the linear effect of age on six morphometric features. Globally, the three macro-structural MRI metrics (GM, CT, SA), decreased over the course of adolescence (GM: $t = -5.6, P_{FDR} < 0.05$; CT: $t = -5.1, P_{FDR} < 0.05$; SA: $t = -3.6, P_{FDR} < 0.05$). The T1w/T2w-ratio increased ($t = 4.0, P_{FDR} < 0.05$), MD showed no significant changes ($t = -1.5, P = 0.13$) and FA decreased ($t = -2.5, P_{FDR} < 0.05$) (**SI Fig. 16B**). Next, we estimated the linear effect of age on six morphometric features at each of 358 cortical areas to resolve the regional anatomical patterning of developmental changes in macro- and micro-structural MRI metrics during adolescence. We generally observed decreases ($t < 0$) in macro-structural features and increases ($t > 0$) in

T1w/T2w ratio, with mixed results in FA and MD (**SI Fig. 16**C). These adolescent changes in individual morphometric features from the HCP-D dataset were correlated with the original results from the NSPN sample as follows: FA: $\rho = 0.03, P_{spin} = 0.12$; MD: $\rho = 0.3, P_{spin} < 0.001$; MT/T1w/T2w-ratio: $\rho = 0.25, P_{spin} < 0.05$; SA: $\rho = 0.26, P_{spin} < 0.001$; GM: $\rho = 0.52, P_{spin} < 0.001$; CT: $\rho = 0.34, P_{spin} < 0.001$. We then estimated linear changes in morphometric similarity with age at each region. These changes were significant in 7 regions after correction for multiple comparisons at $P_{FDR} < 0.05$ (**SI Fig. 17A**). We estimated the mean effect of age on all regions within each of the Mesulam cytoarchitectonic zones Mesulam 1998; Mesulam 2000 and found that morphometric similarity increased in paralimbic regions and decreased in idiotypic and isocortical zones (**SI Fig. 17B**). Lastly, we correlated the effect of age on MSN degree in the HCP-D sample with the effect of age on MSN degree in the NSPN sample and found a significant positive correlation between the two ($\rho = 0.5, P_{spin} < 0.0001$; **SI Fig. 17C**).

#### Limitations of independent replication analysis

We note that the replication of our results in the HCP-D sample may have been limited by a number of factors, including in particular the cross-sectional nature of the sample, the differing age-range, the fact that no MT data were available (requiring approximate substitution as a proxy for intracortical myelination using the T1w/T2w ratio), and differing processing pipelines. However, while we observe some discrepancies in the adolescent development of individual morphometric features between the datasetss, age-related changes in degree of morphometric similarity were remarkably similar in both HCP-D and NSPN samples.

#### Rank-stability of morphometric similarity networks

Rank-stability of MSN global and regional degree: We assessed the longitudinal stability of MSNs between visits for all subjects that had both a baseline and one year follow up visit. Specifically, we ranked subjects by their global and regional MSN degree respectively, and estimated the correlation between the ranked vectors. This provides an estimate of the MSN longitudinal stability, i.e. a positive correlation between baseline and follow up ranks suggests that subjects that had stronger morphometric similarity compared to other subjects at baseline, tended to have a strong MSN degree at follow up as well. Overall, we observe positive relationships between subjects' global and regional ranks, with 241 regions showing a significant correlation between baseline and follow up ranks (**SI Fig. S24B**). A number of factors will influence this analysis: first of all, we observed decreasing macrostructural and increasing microstructural feature strength. The crossing patterns of development of these individual features will inevitably affect morphometric similarity. Secondly, the accelerated longitudinal design of the NSPN study means that while subjects were uniformly assessed with a one year gap between baseline and follow up scans, the baseline scans were acquired between the ages of 14 and 25. It is conceivable that the changes in morphometric similarity are stronger or weaker at different periods between these two ages, which may affect the rank-reordering of subjects.

### NSPN Consortium Author List

**Principal investigators:**
Edward Bullmore (CI from 01/01/2017)
Raymond Dolan
Ian Goodyer (CI until 01/01/2017)
Peter Fonagy
Peter Jones

**NSPN (funded) staff:**
Michael Moutoussis
Tobias Hauser
Sharon Neufeld
Rafael Romero-García
Michelle St Clair
Petra Vértes
Kirstie Whitaker
Becky Inkster
Gita Prabhu
Cinly Ooi
Umar Toseeb
Barry Widmer
Junaid Bhatti
Laura Villis
Ayesha Alrumaithi
Sarah Birt
Aislinn Bowler
Kalia Cleridou
Hina Dadabhoy
Emma Davies
Ashlyn Firkins
Sian Granville
Elizabeth Harding
Alexandra Hopkins
Daniel Isaacs
Janchai King
Danae Kokorikou
Christina Maurice
Cleo McIntosh
Jessica Memarzia
Harriet Mills
Ciara O'Donnell
Sara Pantaleone
Jenny Scott

**Affiliated scientists:**
Pasco Fearon
John Suckling
Anne-Laura van Harmelen
Rogier Kievit

| agebin | Baseline female | Baseline male | 6 Months female | 6 Months male | Follow Up female | Follow Up male |
|---|---|---|---|---|---|---|
| 1 | 29 | 28 | 3 | 2 | 19 | 25 |
| 2 | 34 | 28 | 2 | 2 | 24 | 22 |
| 3 | 23 | 24 | 3 | 2 | 14 | 13 |
| 4 | 30 | 32 | 5 | 5 | 15 | 21 |
| 5 | 20 | 18 | 1 | 2 | 8 | 15 |

**Table 1: NSPN structural MRI data sample overview:** The NSPN sample was a sex-balanced, age-stratified longitudinal cohort, with subjects recruited in five age bins: 14-15 years, 16-17 years, 18-19 years, 20-21 years and older than 22 years at baseline. Approximately 30 subjects per sex were recruited in each age bin. Subjects were invited for scanning at a baseline and follow-up visit approximately one year later, with a small subset of subjects also invited for an intermediate scan about six months after baseline. Here, we list the number of structural scans available in the final sample (after quality control) per age-bin, for each of the visits, stratified by sex.

| | session | sex | sMRI available | fMRI available |
|---|---|---|---|---|
| 1 | ses-baseline | female | 136 | 132 |
| 2 | ses-baseline | male | 130 | 122 |
| 3 | ses-6Month | female | 14 | 14 |
| 4 | ses-6Month | male | 13 | 12 |
| 5 | ses-1stFollowUp | female | 80 | 78 |
| 6 | ses-1stFollowUp | male | 96 | 90 |

**Table 2: Available data:** Here, we list the number of scans per session for sMRI and fMRI data.

| | $t_{age}$ | $P_{age}$ | $t_{sex}$ | $P_{sex}$ | $P_{age_{FDR}}$ | $P_{sex_{FDR}}$ |
|---|---|---|---|---|---|---|
| Fractional Anisotropy | 1.78 | 0.08 | 1.13 | 0.26 | 0.09 | 0.31 |
| Mean Diffusivity | -0.42 | 0.67 | -2.01 | 0.05 | 0.67 | 0.07 |
| Magnetization Transfer | 3.19 | 0.00 | -2.10 | 0.04 | 0.00 | 0.07 |
| Surface Area | -2.33 | 0.02 | 2.20 | 0.03 | 0.03 | 0.07 |
| Grey Matter Volume | -5.23 | 0.00 | 2.85 | 0.00 | 0.00 | 0.03 |
| Cortical Thickness | -7.29 | 0.00 | -0.11 | 0.91 | 0.00 | 0.91 |

**Table 3: Age and sex effects on individual morphometric features:** We estimated the linear effect of age on individual morphometric featurea (FA, MD, MT, SA, GM, CT) using linear mixed effects models with a fixed effect of age, sex and site, and a random effect of subject. Above, we list the $t$ and $P$-values from this model.

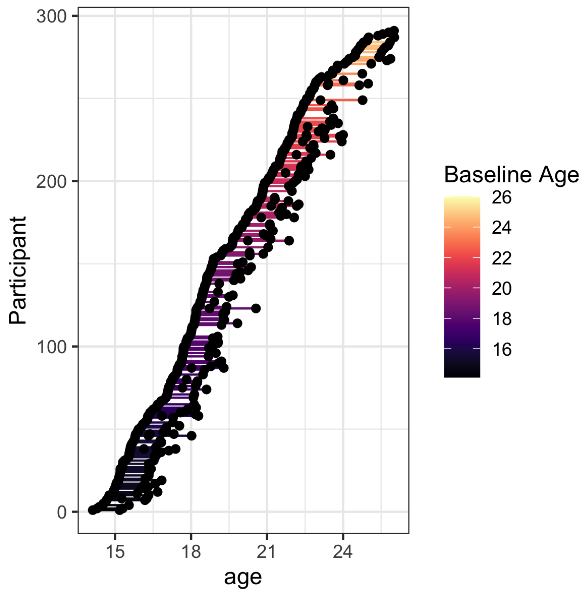

**Fig. 1: Acquisition overview:** The NSPN study was acquired as an accelerated longitudinal study, with participants recruited in five age bins (14-15 years, 16-17 years, 18-19 years, 20-21 years, 22 years and older), stratified by sex. Here, we illustrate the study design by showing the each of each participant at their respective baseline, 6 month and 1 year follow up scan.

| Feature | Zone 1 | Zone 2 | $P$ | $P_{FDR}$ | Significance |
|---|---|---|---|---|---|
| FA | idiotypic | unimodal | 0.3260962 | 6.5e-01 | ns |
| | idiotypic | paralimbic | 0.0000079 | 2.4e-05 | **** |
| | idiotypic | heteromodal | 0.0000002 | 8.0e-07 | **** |
| | unimodal | paralimbic | 0.0000049 | 1.9e-05 | **** |
| | unimodal | heteromodal | 0.0000000 | 0.0e+00 | **** |
| | paralimbic | heteromodal | 0.9171796 | 9.2e-01 | ns |
| MD | idiotypic | unimodal | 0.0816691 | 2.4e-01 | ns |
| | idiotypic | paralimbic | 0.0016081 | 6.4e-03 | ** |
| | idiotypic | heteromodal | 0.1221331 | 2.4e-01 | ns |
| | unimodal | paralimbic | 0.0800181 | 2.4e-01 | ns |
| | unimodal | heteromodal | 0.0000229 | 1.1e-04 | **** |
| | paralimbic | heteromodal | 0.0000004 | 2.7e-06 | **** |
| MT | idiotypic | unimodal | 0.0002239 | 1.3e-03 | *** |
| | idiotypic | paralimbic | 0.3247295 | 3.2e-01 | ns |
| | idiotypic | heteromodal | 0.0227680 | 9.1e-02 | * |
| | unimodal | paralimbic | 0.0008349 | 4.2e-03 | *** |
| | unimodal | heteromodal | 0.0228688 | 9.1e-02 | * |
| | paralimbic | heteromodal | 0.1303646 | 2.6e-01 | ns |
| SA | idiotypic | unimodal | 0.0399473 | 8.0e-02 | * |
| | idiotypic | paralimbic | 0.3186235 | 3.2e-01 | ns |
| | idiotypic | heteromodal | 0.0000359 | 1.8e-04 | **** |
| | unimodal | paralimbic | 0.0002117 | 8.5e-04 | *** |
| | unimodal | heteromodal | 0.0033420 | 1.0e-02 | ** |
| | paralimbic | heteromodal | 0.0000000 | 0.0e+00 | **** |
| GM | idiotypic | unimodal | 0.7056571 | 7.1e-01 | ns |
| | idiotypic | paralimbic | 0.0072423 | 1.4e-02 | ** |
| | idiotypic | heteromodal | 0.0000001 | 5.0e-07 | **** |
| | unimodal | paralimbic | 0.0003584 | 1.1e-03 | *** |
| | unimodal | heteromodal | 0.0000000 | 0.0e+00 | **** |
| | paralimbic | heteromodal | 0.0000000 | 0.0e+00 | **** |
| CT | idiotypic | unimodal | 0.3915979 | 3.9e-01 | ns |
| | idiotypic | paralimbic | 0.0000529 | 1.4e-04 | **** |
| | idiotypic | heteromodal | 0.0000473 | 1.4e-04 | **** |
| | unimodal | paralimbic | 0.0000000 | 0.0e+00 | **** |
| | unimodal | heteromodal | 0.0000162 | 6.5e-05 | **** |
| | paralimbic | heteromodal | 0.0000000 | 0.0e+00 | **** |

**Table 4: Pairwise comparison of mean changes in individual morphometric features by Mesulam class: We used a Wilcoxon test to estimated the difference in the mean rate of change between Mesulam cytoarchiteconic classes within each morphometric feature. See also SI Fig. 7**

| region | $t$-value | $P_{FDR}$ | mesulam class |
|---|---|---|---|
| L POS2 | -3.02 | 0.04 | heteromodal |
| L IP2 | -3.38 | 0.02 | heteromodal |
| R RSC | -4.02 | 0.00 | heteromodal |
| R 7PC | -4.09 | 0.00 | heteromodal |
| R IFJp | -2.93 | 0.04 | heteromodal |
| R 46 | -2.94 | 0.04 | heteromodal |
| R 10v | -3.13 | 0.03 | heteromodal |
| R PFt | -4.00 | 0.00 | heteromodal |
| R DVT | -3.15 | 0.03 | heteromodal |
| R TE1m | 4.52 | 0.00 | heteromodal |
| L 6r | -3.27 | 0.03 | idiotypic |
| R V8 | -2.93 | 0.04 | idiotypic |
| L p24pr | 3.02 | 0.04 | paralimbic |
| L a24pr | 3.10 | 0.03 | paralimbic |
| L a24 | 3.20 | 0.03 | paralimbic |
| L 47s | 2.96 | 0.04 | paralimbic |
| L PoI2 | 4.34 | 0.00 | paralimbic |
| L MI | 3.49 | 0.02 | paralimbic |
| L PeEc | 2.90 | 0.05 | paralimbic |
| L VVC | 3.94 | 0.00 | paralimbic |
| L pOFC | 3.09 | 0.03 | paralimbic |
| L a32pr | 4.57 | 0.00 | paralimbic |
| R 33pr | 3.74 | 0.01 | paralimbic |
| R p32pr | 3.69 | 0.01 | paralimbic |
| R PFcm | 4.29 | 0.00 | paralimbic |
| R FOP4 | 5.04 | 0.00 | paralimbic |
| R H | 3.27 | 0.03 | paralimbic |
| R 31a | 3.93 | 0.00 | paralimbic |
| R s32 | 3.45 | 0.02 | paralimbic |
| R PI | 3.17 | 0.03 | paralimbic |
| L STGa | -3.07 | 0.04 | unimodal |
| R 1 | -3.02 | 0.04 | unimodal |
| R TPOJ2 | -3.15 | 0.03 | unimodal |

**Table 5: Significant regional effects of age on morphometric similarity:** We estimated the linear effect of age on morphometric similarity using linear mixed effects models with a fixed effect of age, sex and site, and a random effect of subject. We find that 33 regions show significant effects of age after FDR-correction. Above, we list the $t$ and $P$-values for these regions, together with their respective Mesulam (Mesulam 2000; Mesulam 1998) class assignment.

| Zone 1 | Zone 2 | $P$ | $P_{FDR}$ | Significance | Method |
|---|---|---|---|---|---|
| heteromodal | idiotypic | 0.78 | 0.78 | ns | Wilcoxon |
| heteromodal | paralimbic | 0.00 | 0.00 | **** | Wilcoxon |
| heteromodal | unimodal | 0.00 | 0.00 | *** | Wilcoxon |
| idiotypic | paralimbic | 0.00 | 0.00 | **** | Wilcoxon |
| idiotypic | unimodal | 0.00 | 0.01 | ** | Wilcoxon |
| paralimbic | unimodal | 0.00 | 0.00 | **** | Wilcoxon |

**Table 6: Pairwise comparison of mean changes morphometric similarity by Mesulam class**: We used a Wilcoxon test to estimated the difference in the mean rate of change in morphometric similarity between Mesulam cytoarchiteconic classes. See also **Fig. 2** in the main text.

| Parameter of Change | Zone 1 | Zone 2 | $P$ | $P_{FDR}$ | Significance |
|---|---|---|---|---|---|
| Baseline Coupling | heteromodal | idiotypic | 0.0003537 | 1.4e-03 | *** |
| | heteromodal | paralimbic | 0.0000005 | 3.1e-06 | **** |
| | heteromodal | unimodal | 0.2415052 | 4.8e-01 | ns |
| | idiotypic | paralimbic | 0.5412466 | 5.4e-01 | ns |
| | idiotypic | unimodal | 0.0117424 | 3.5e-02 | * |
| | paralimbic | unimodal | 0.0002896 | 1.4e-03 | *** |
| Rate of Change in Coupling | heteromodal | idiotypic | 0.9897100 | 9.9e-01 | ns |
| | heteromodal | paralimbic | 0.0081024 | 4.1e-02 | ** |
| | heteromodal | unimodal | 0.1481998 | 4.4e-01 | ns |
| | idiotypic | paralimbic | 0.0429494 | 1.7e-01 | * |
| | idiotypic | unimodal | 0.4110969 | 8.2e-01 | ns |
| | paralimbic | unimodal | 0.0004275 | 2.6e-03 | *** |

**Table 7: Pairwise comparison of adolescent parameters of development in structure-function coupling by Mesulam class:** We used a Wilcoxon test to estimated the difference in the mean rate of change in morphometric similarity between Mesulam cytoarchiteconic classes. See also **SI Fig. 19**

| Zone 1 | Zone 2 | $P$ | $P_{FDR}$ | Significance |
|---|---|---|---|---|
| idiotypic | unimodal | 0.79 | 1.00 | ns |
| idiotypic | paralimbic | 0.02 | 0.08 | * |
| idiotypic | heteromodal | 0.99 | 1.00 | ns |
| unimodal | paralimbic | 0.00 | 0.03 | ** |
| unimodal | heteromodal | 0.89 | 1.00 | ns |
| paralimbic | heteromodal | 0.01 | 0.05 | * |

**Table 8: Pairwise comparison of adolescent changes in participation coefficient by Mesulam zone:** We used a Wilcoxon test to estimated the difference in the mean rate of change in participation coefficient between Mesulam cytoarchiteconic zones. See also **SI Fig. 23**

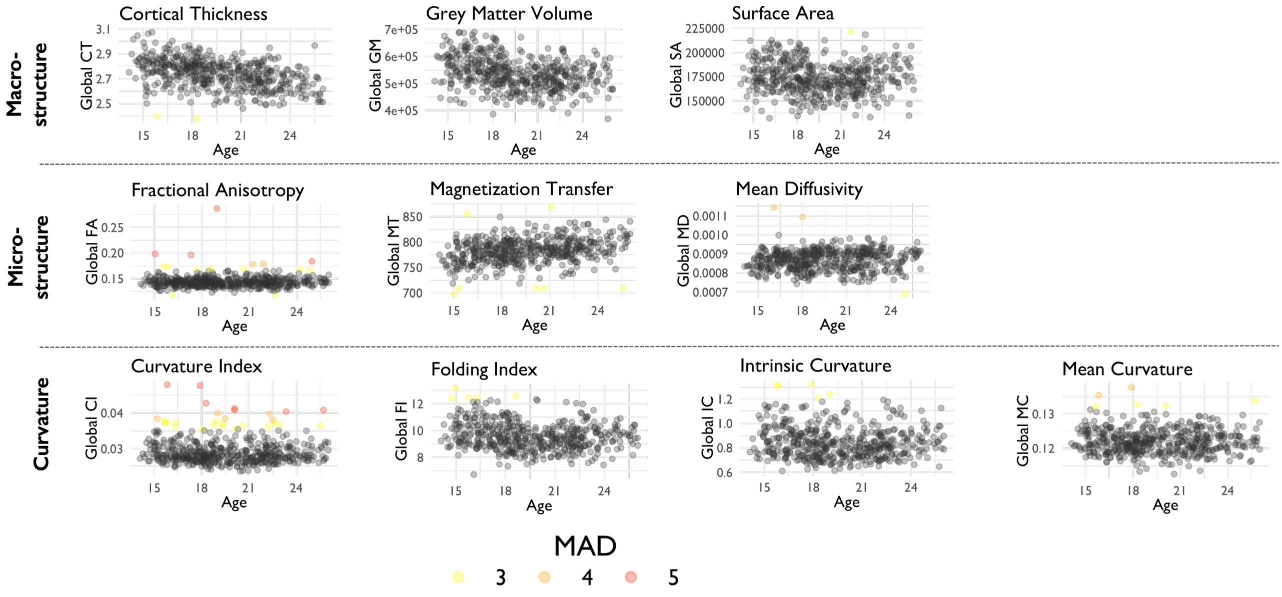

**Fig. 2: Global morphometric outliers:** We estimated MAD scores across subjects for each of 10 morphometric features: CT, GM, SA, FA, MT, MD, IC, FI, CI, MC. We defined outliers as subjects with $MAD \geq 5$ in at least one morphometric feature. Here, global (raw) feature values, colored by their respective MAD scores, are shown to highlight which datapoints were removed.

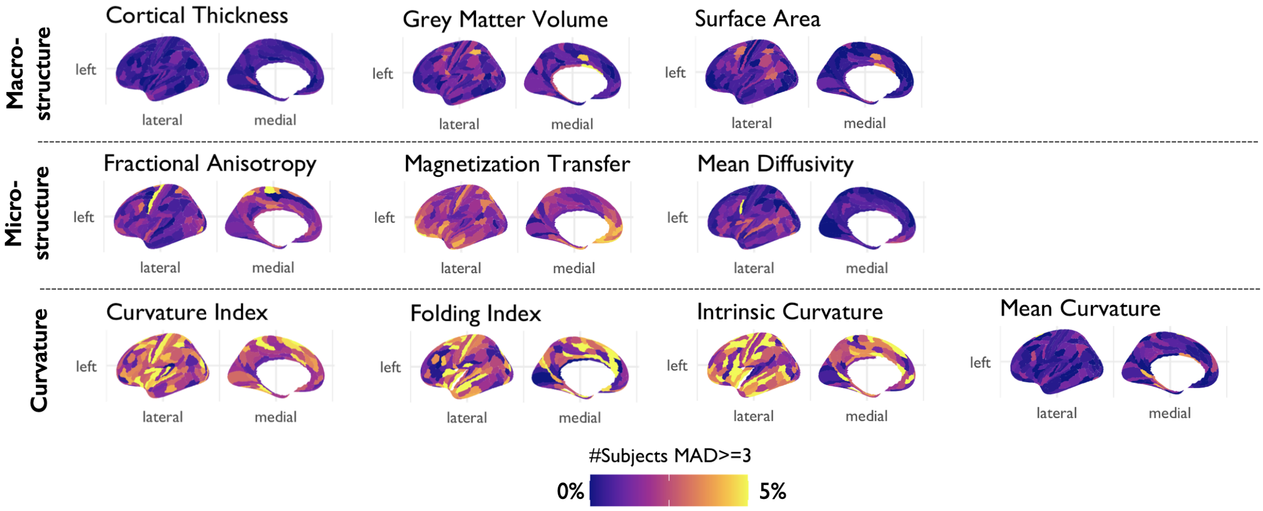

**Fig. 3: Local morphometric outliers:** We estimated the local MAD score for each subject at each region within each morphometric feature. Here, we show the percentage of subjects with $MAD \geq 5$ in each region. Due to elevated rates of outliers compared to other measures, the curvature features were excluded from further analysis.

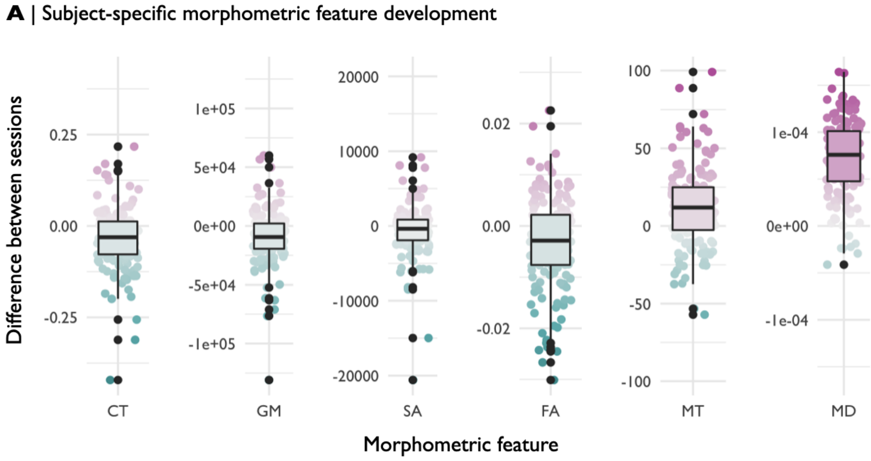

**Fig. 4: Subject specific global differences in morphometric features between visits:** For the subset of subjects that had a baseline and follow up visit ($N = 171$), we estimated the global difference in morphometric feature strength between visits. We find that overall, the subject-specific difference in feature strength between visits is associated with the between-subject trend, i.e. global MT increases with age across subjects, and on the subject-specific level, the majority of subjects showed increased global MT at follow up compared to baseline. This trend is true for all features except fractional anisotropy, which shows non-significant ($P_{FDR} >= 0.05$) increases with age across subjects, but on the subject-specific level, the majority of subjects have decreased global FA at follow up. We further assessed the percentage of subject that follows the global trend when assessing the raw difference in their global feature values at baseline compared to follow up and find the following: GM: $71\%$; CT: $69\%$; SA: $58\%$; FA: $39\%$; MT $71\%$; MD: $95\%$

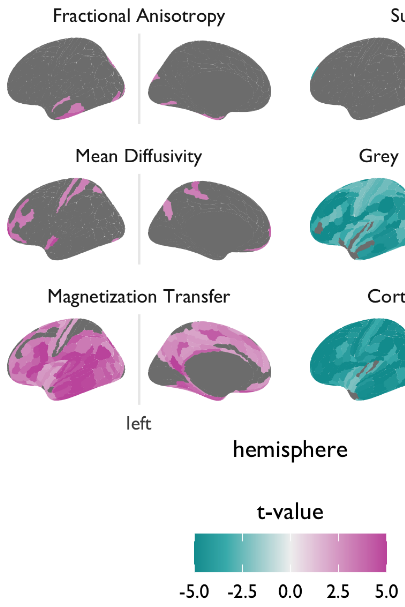

**Fig. 5: Age-related changes in individual morphometric features:** We modeled the linear effect of age on six morphometric features at each of 358 cortical areas to resolve the anatomical patterning of decreased macro- and increased micro-structural metrics during adolescence. We largely observed increases ($t > 0$) in micro-structural features, and decreases ($t < 0$) in macro-structural features. (B) We thresholded the results for significance after correction for multiple comparisons, $P_{FDR} < 0.05$. The results were highly symmetric across hemispheres, so here only the left hemisphere is shown.

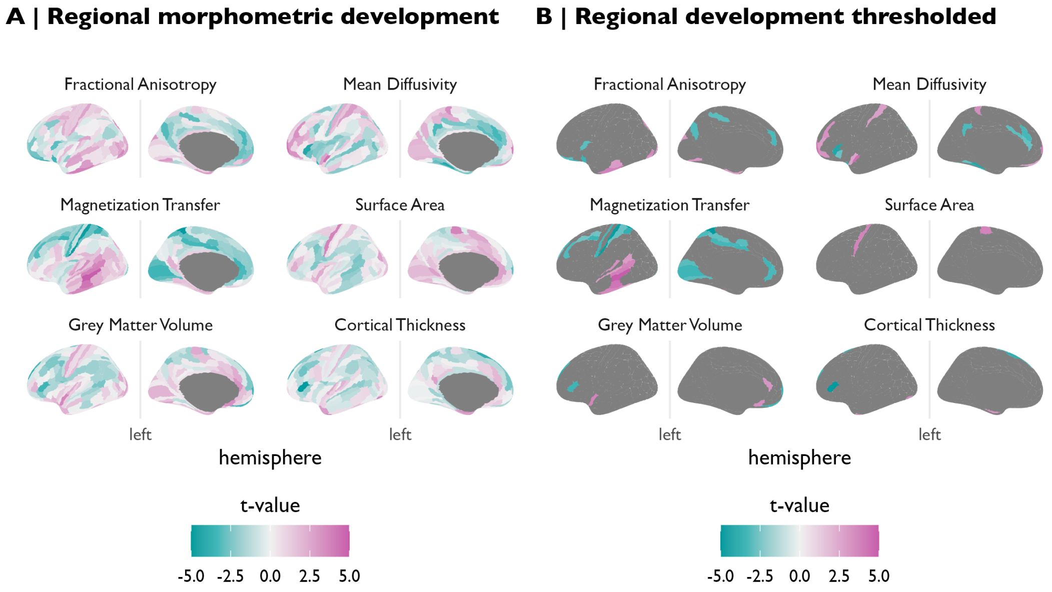

**Fig. 6: Adolescent changes in macro-structural and micro-structural MRI metrics corrected for global effects of age:** We estimated age-related changes in regional features correcting for their respective global values, i.e., regional rates of change relative to each feature's global rate of change. (A) We modeled the linear effect of age on 6 micro-structural and macro-structural MRI features in each of 358 cortical areas to resolve the anatomical patterning of decreased macro- and increased micro-structural metrics during adolescence. (B) We thresholded the results from panel (A) for significance after correction for multiple comparisons $P_{FDR} < 0.05$. The results were highly symmetric across hemispheres, so here only the left hemisphere is shown.

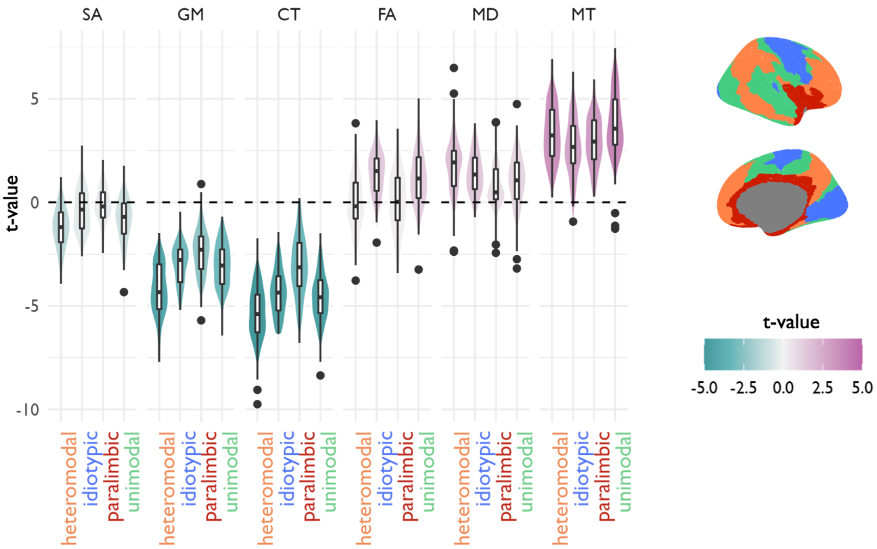

**Fig. 7: Age-related changes in individual morphometric features by mesulam classes:** Here, we summarize the linear effect of age on six morphometric features at each of 358 cortical areas by Mesulam classes. Overall, we observe increases $(t > 0)$ in micro-structural features, and decreases $(t < 0)$ in macro-structural features. More specifically, we found that in macro-structural metrics, paralimbic areas tended to decrease less than mesocortical areas. Conversely, in micro-structural metrics, paralimbic areas tended to increase less than mesocortical areas. See **SI Table 4** for pairwise comparison between Mesulam zones by morphometric feature.

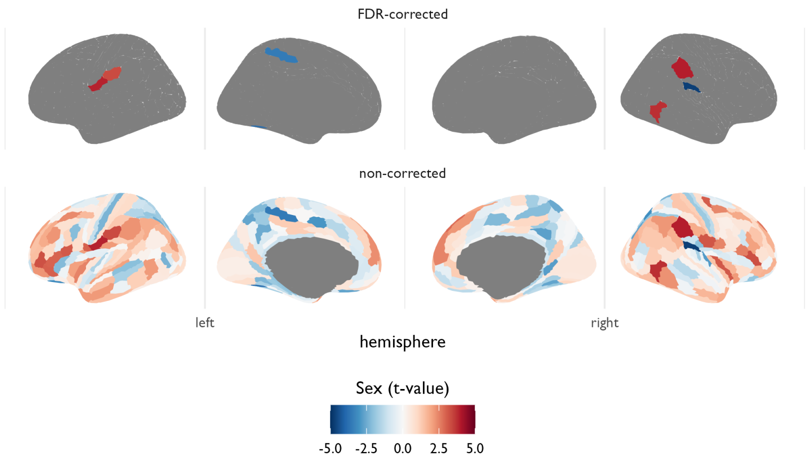

**Fig. 8: Sex effects on adolescent changes in morphometric similarity:** We estimated sex effects on adolescent changes in morphometric similarity at each region (*bottom*). Positive $t$-values indicate that morphometric similarity increased with age more strongly in males compared to females. After correction for multiple comparisons, we found that seven regions (*L_5mv, L_OP4, L_IP0, L_VMV1, R_52, R_TF, R_IP0*) displayed significant sex differences in age-related changes in morphometric similarity ($P_{FDR} < 0.05$). Overall, the pattern of observed (non-significant) sex effects included increased morphometric similarity in females in limbic and default mode network regions, and increased morphometric similarity in males elsewhere in the cortex.

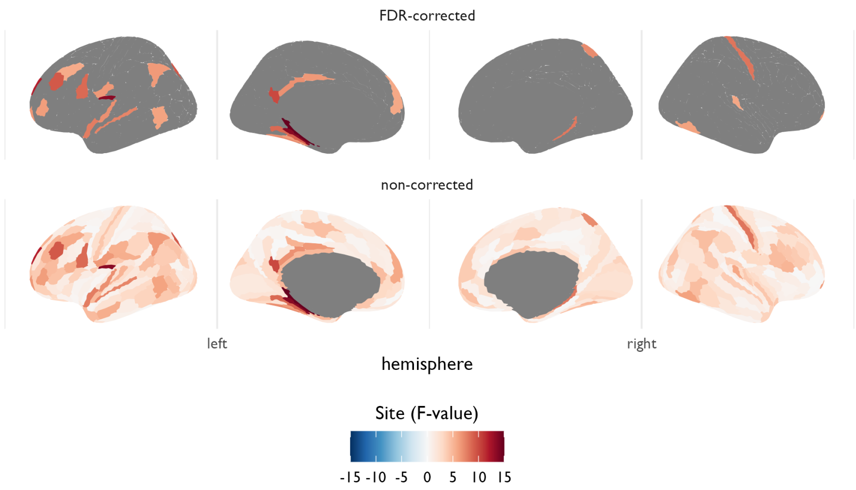

**Fig. 9: Site effects on adolescent changes in morphometric similarity:** The MRI data used for this work was acquired at three different sites (Kiddle et al. 2017). We estimated age effects on regional morphometric similarity degree, $k$, using linear mixed effects models with a fixed effect of age, sex and site, and a random effect of subject. Here, we show the effect of site ($F$-value) from those models (*bottom*). We find that 30 regions showed significant site effects, however only two regions showed both significant age-related changes in MSN weighted degree and effects of site (*L_6r, R_10v*), thus our strongest effects should be largely unaffected by site differences.

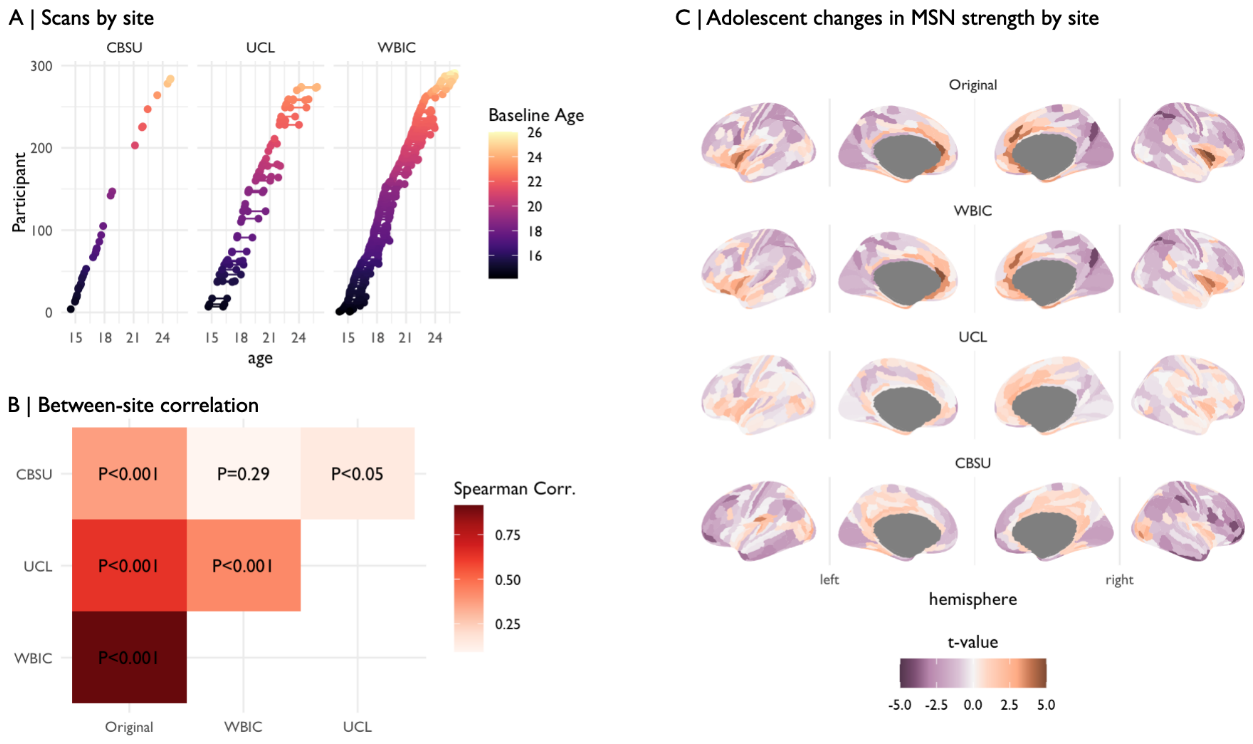

**Fig. 10: Changes in morphometric similarity by scanning site:** The MRI data used for this work was acquired at three different sites (Kiddle et al. 2017). In order to assess the sensitivity of our main results to the effects of scanning we split the full sample by site and independently estimated the effects of age on adolescent development of morphometric similarity. (A) Overview of the number of scans acquired at each site: Wolfson Brain Imaging Center Cambridge (WBIC, N = 347); University College London (UCL; N = 98); Cambridge MRC Cognition and Brain Sciences Unit, Cambridge (CBSU; N = 33). (B) Using Spearman's correlation ($\rho$), we find that the site-specific maps of adolescent changes in morphometric similarity are significantly positively ($P_{spin} < 0.001$) correlated with the original map. (C) Visualization of the effect of age on morphometric similarity by site.

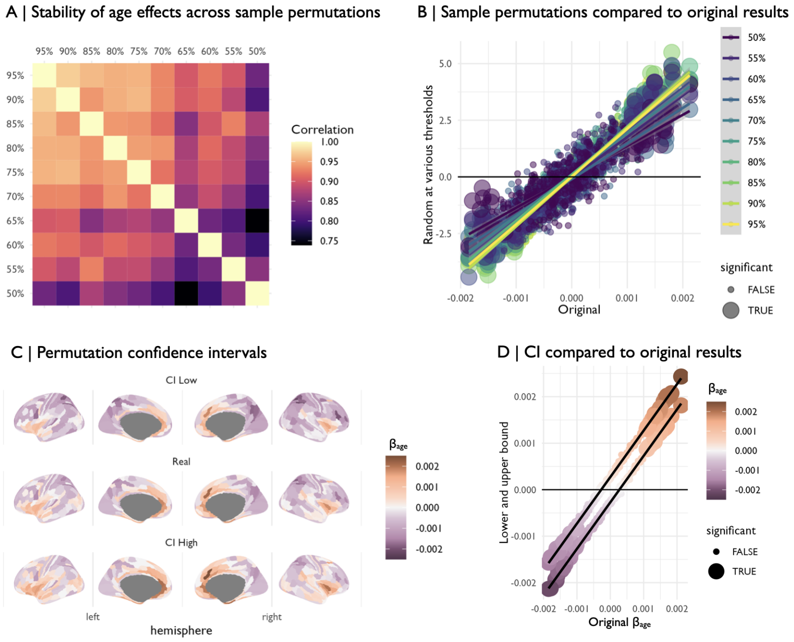

**Fig. 11: Stability of age effects on morphometric similarity:** We tested for the sensitivity of the age effects on morphometric similarity to the specific sample used. To this end, we randomly selected subsets of the data, maintaining the age- and sex-stratified nature of the sample. (A) First, we randomly selected decreasing percentages of subjects in 5% steps and tested for age effects on regional morphometric similarity as in the main analysis in these smaller samples. We then correlated the $t$-statistics of age estimated in this ablation analysis with the original $t$-statistic from the main analysis. We find a high spatial correlation between the original map and the maps derived from the random subsets of the data. As expected, the correlation to other maps decreases with decreasing sample sizes of the random subsets, but the correlation stays high ($r > 0.75$). (B) Importantly, the none of the regions found to show significant changes in morphometric similarity after correction for multiple comparisons (large dots in the plot) in the original analysis changed sign in any of the random samples. (C) Next, we derived lower and upper permutation-based confidence for visual comparison of the robustness of the directionality of observed age effects on morphometric similarity. To this end, we performed a leave-N-out-analysis at a single threshold of 10%. More specifically, in 1000 permutations we randomly dropped 10% of subjects for each sex and agebin and re-estimated the $t$-statistic of age on MSN weighted degree. Within each region, we then estimated the standard deviation (SD) of the permutation distribution and derived regional upper and lower bounds at $1.96 * SD$ around the original $\beta$-coefficient of age. We estimated these confidence intervals around the $\beta$-coefficient of age rather than the $t$-statistic, since the random subsets are estimated on a smaller sample size than the original sample, thus $t$ values are not directly comparable. We find that the lower and upper boundary maps look almost identical to the original map. (D) Again, none of the regions with significant changes in MSN weighted degree in the original sample (marked as large dots in the plot) changed sign in the upper or lower confidence bounds.

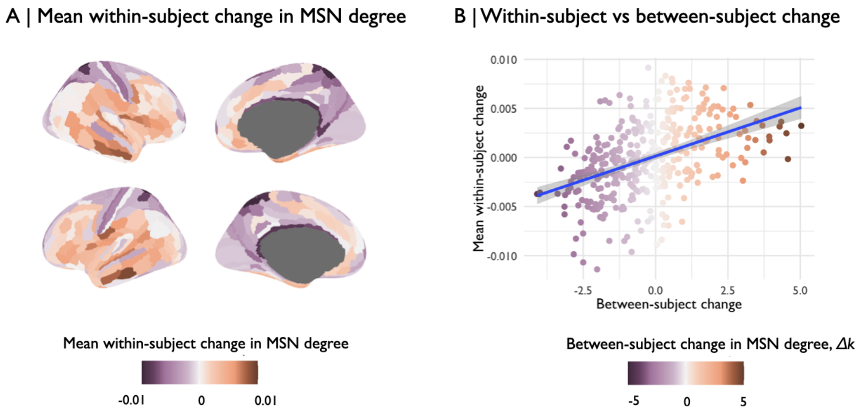

**Fig. 12: Within vs. between-subject regional changes in MSN degree:** The longitudinal nature of the study allows us to contrast the reported between-subject change in MSN degree ($\Delta k$) with within-subject changes in regional MSN development. We opted to estimate the raw within-subject difference in regional MSN degree for subjects that had both a baseline and one year follow-up scan for contrasting with the group-level results. (A) We estimated the mean within-subject change in MSN degree at each region. (B) We find that the mean within-subject change in MSN degree is significantly positively ($\rho = 0.41, P_{spin} < 0.001$) correlated with the between-subject rate of change. We thus conclude that there is reasonable correspondence between between-subject and within-subject effects in MSN development.

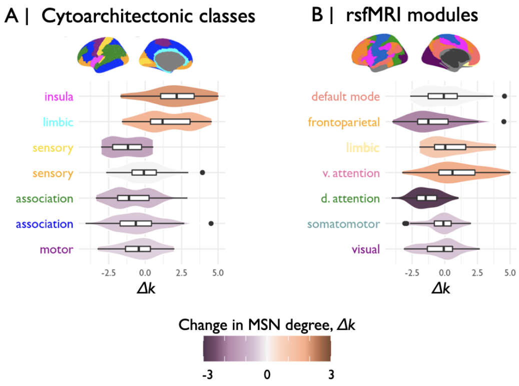

**Fig. 13: Adolescent rate of change in MSN degree by cytoarchitectonic and functional classes:** (A) We estimated the mean effect of age on morphometric similarity in all regions within each of the von Economo cytoarchitectonic classes and found that morphometric similarity increased in insular and limbic cytoarchitectonic classes and decreased in all other classes. (B) We further estimated the mean effect of age on all regions within each pre-defined fMRI modules (Yeo et al. 2011) and found that morphometric similarity increased in limbic and ventral attention functional networks and decreased most strongly in dorsal attention and frontoparietal networks.

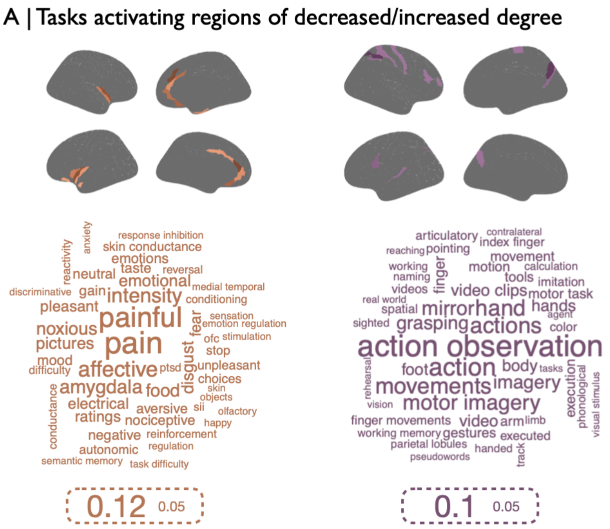

**Fig. 14:** Word cloud of cognitive terms (Yarkoni et al. 2011) scaled according to their strength of association with the thresholded ($P_{FDR} < 0.05$) map of adolescent increases ($t > 0$, orange) and decreases ($t < 0$, purple) in morphometric similarity.

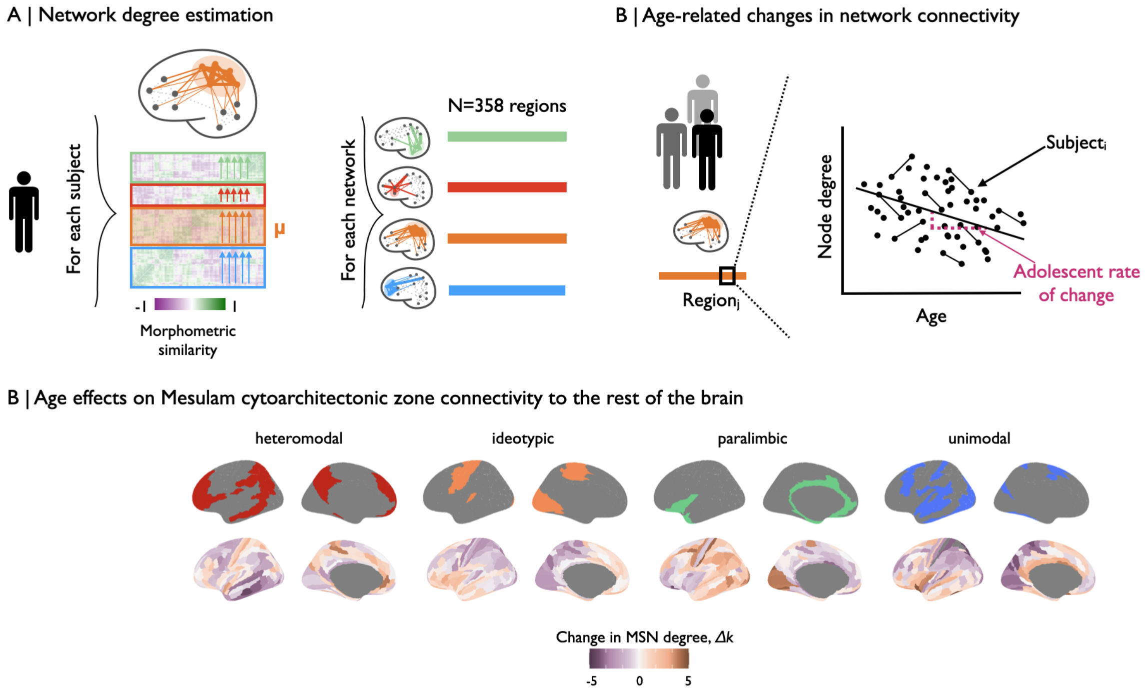

**Fig. 15: Cytoarchitectonic class-specific changes in morphometric similarity:** (A) For each subject, within each cytoarchitectonic zone (Mesulam 1998; Mesulam 2000), we estimated the strength of similarity of all other cortical regions to the nodes in a given zone by summing the relevant sections of the connectivity matrix. Thus within each subject, for a given cytoarchitectonic zone with $Z$ regions, we averaged the connectivity of these $Z$ regions to all $N = 358$ regions included in our analyses. For each subject, this resulted in a $1 \times N = 358$ connectivity vector for each Mesulam zone which summarised the connectivity of regions in a given zone to the rest of the brain. (B) Then, we then estimated age-related changes in morphometric similarity in each of $N = 358$ regions for each Mesulam class separately. We thus derived the rate of change in morphometric similarity of the connectivity between a given Mesulam zone and the rest of the brain. (C) We found that the patterns of adolescent rate of change in morphometric similarity differed qualitatively between cytoarchitectonic zones.

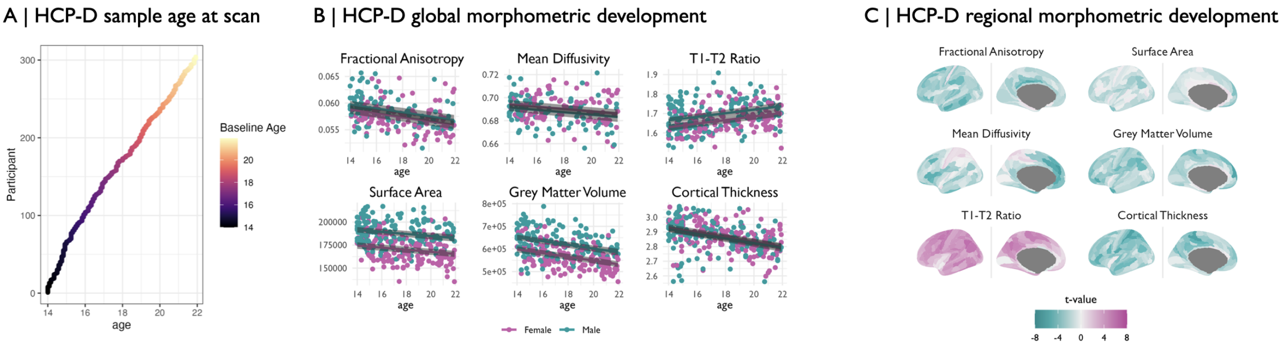

**Fig. 16: Global and regional morphometric development in the HCP-D sample:** (A) The HCP-D sample is a cross sectional cohort of children and adolescents aged 5 to 21 years. For better comparison with our primary cohort, we included only subjects aged 14 and older. (B) We modelled the linear effect of age on six morphometric features in the HCP-D sample. Globally, the three macro-structural MRI metrics (GM, CT, SA), decreased over the course of adolescence (GM: $t = -5.6, P_{FDR} < 0.05$; CT: $t = -5.1, P_{FDR} < 0.05$; SA: $t = -3.6, P_{FDR} < 0.05$). The T1w/T2w-ratio increased ($t = 4.0, P_{FDR} < 0.05$), MD showed no significant changes ($t = -1.5, P = 0.13$) and FA decreased ($t = -2.5, P_{FDR} < 0.05$) (**SI Fig. 16B**). (C) We modelled the linear effect of age on six morphometric features at each of 358 cortical areas to resolve the anatomical patterning of developmental changes in macro- and micro-structural MRI metrics during adolescence. We generally observed decreases ($t < 0$) in macro-structural features and increases ($t > 0$) in T1w/T2w ratio, with mixed results in FA and MD.

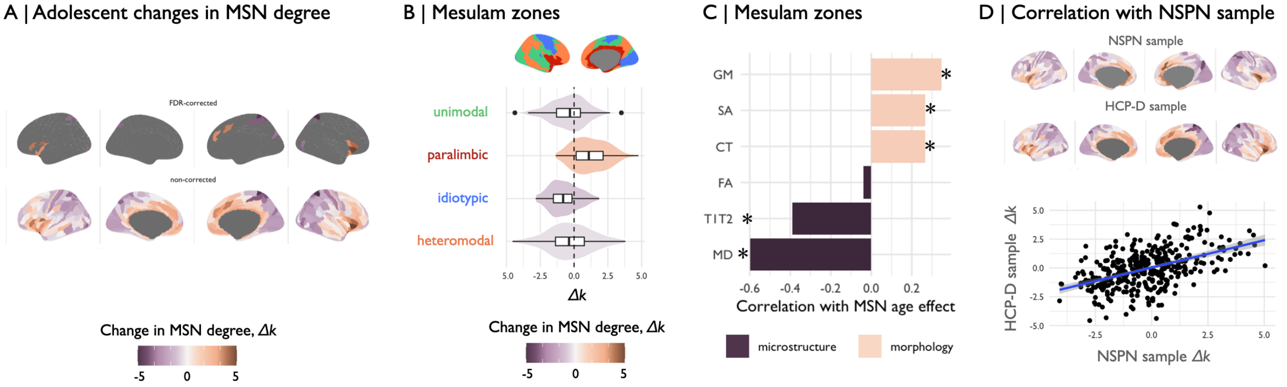

**Fig. 17: Adolescent change in degree of morphometric similarity, $\Delta k$, in the HCP-D sample:** We estimated morphometric similarity networks for each subject by correlating the standardized morphometric feature vectors for each possible pair of regions. (A) We estimated linear changes in morphometric similarity with age, $\Delta k$, at each region and found that morphometric similarity decreased in neocortical (frontal, occipital) regions, and increased in medial and temporal cortical regions. These changes were significant after correction for multiple comparisons in 7 regions. (B) We estimated the mean effect of age on all regions within each of the Mesulam cytoarchitectonic zones Mesulam 1998; Mesulam 2000 and found that on average morphometric similarity increased in paralimbic regions and decreased otherwise. (C) Then, we assessed the correlation between adolescent effects on individual MRI features at each region and the adolescent effect on degree of morphometric similarity, or "hubness", of each regional node in the cortical connectome. We found that the T1w/T2w-ratio and other micro-structural MRI features were negatively correlated with adolescent change in MSN degree, i.e., cortical myelination increased in areas that become more morphometrically dissimilar, or less hub-like with $\Delta k < 0$, during adolescence. Conversely, macro-structural MRI features were positively correlated with adolescent change in MSN degree, i.e., cortical thickness, volume and surface area all decreased in regions that became less hub-like during adolescence. Significant correlations at $P_{spin} < 0.05$ are marked with an asterix. (D) Lastly, we correlated the effect of age on MSN degree in the HCP-D sample with that in the NSPN sample and found a significant positive correlation between the two ($\rho = 0.5, P_{spin} < 0.001$). We thus conclude there is a high spatial correspondance between the effects of age on MSN degree estimated in two fully independent samples.

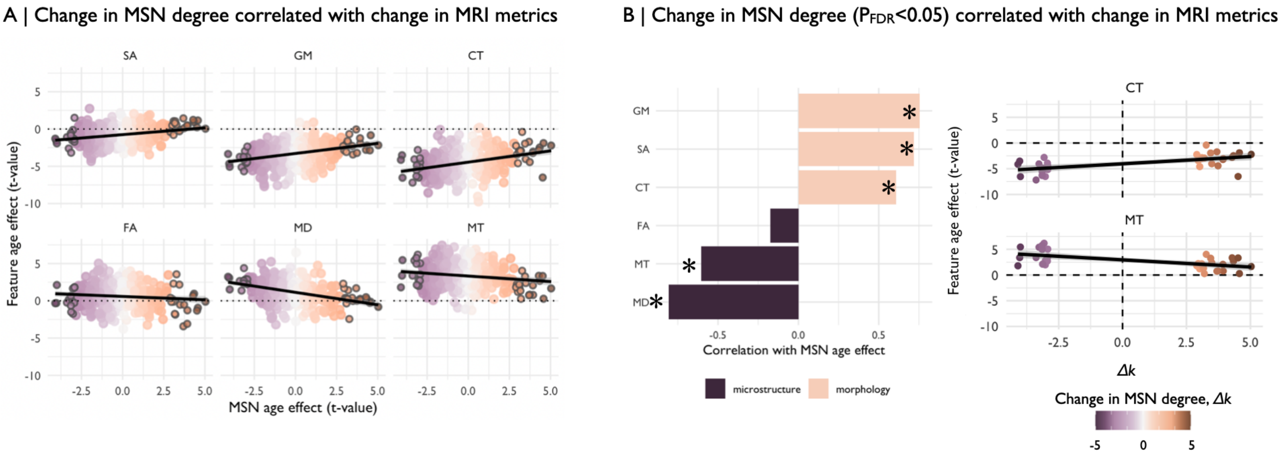

**Fig. 18: Association of adolescent changes in MSN degree with individual MRI metrics:** (A) We estimated the correlation between adolescent effects on individual MRI features at each region and the adolescent effect on degree of morphometric similarity, or "hubness", of each regional node in the cortical connectome. We found that MT and other micro-structural MRI features were negatively correlated with adolescent change in MSN degree, i.e., cortical myelination increased in areas that become more morphometrically dissimilar, or less hub-like with $\Delta k < 0$, during adolescence. Conversely, macro-structural MRI features were positively correlated with adolescent change in MSN degree, i.e., cortical thickness, volume and surface area all decreased in regions that became less hub-like during adolescence. Here, we highlight regions that showed significant changes in morphometric similarity ($P_{FDR} < 0.05$). (B) Additionally, we repeated these analyses using the thresholded map ($P_{FDR} < 0.05$) of adolescent changes in MSN degree only N=33 regions. In line with results presented in **Fig. 3** of the main text and panel (A) of this figure, we find that micro-structural MRI features were negatively correlated with adolescent change in MSN degree, whereas macrostructural measures were positively correlated. Significant correlations ($P_{spin} < 0.05$) are demarcated with an asterix in the left panel.

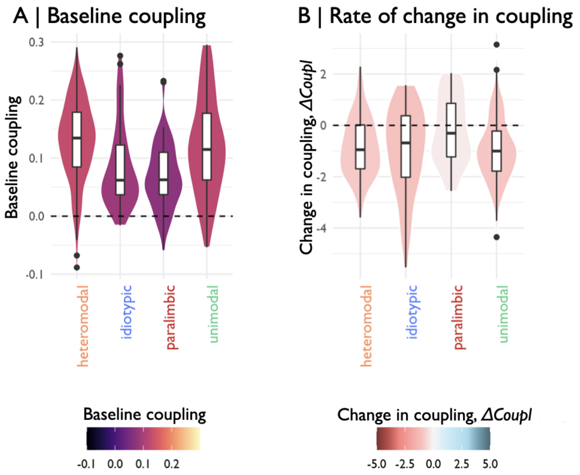

**Fig. 19: Structure-function coupling by cytoarchitectonic zones:** We estimated parameters of adolescent structure-function development by Mesulam zone (Mesulam 1998; Mesulam 2000). Specifically, we estimated the linear change with age in regional structure-function coupling using using linear mixed effects models with a fixed effect of age, sex and site, and a random effect of subject. From this model, we derived the (A) baseline structure-function coupling as the predicted coupling at age 14. We found that baseline structure-function coupling was lower in paralimbic and idiotypic zones, compared to heteromodal and unimodal zones. (B) We further derived the adolescent rate of change in structure-function coupling, as the $t$-value of age from the model. We found that the rate of change in structure-function coupling was more strongly negative ($t < 0$) in iso-cortical areas compared to paralimbic areas, meaning that regions in iso-cortex decoupled more strongly than paralimbic regions. See **SI Table 7** for pairwise comparison between Mesulam zones.

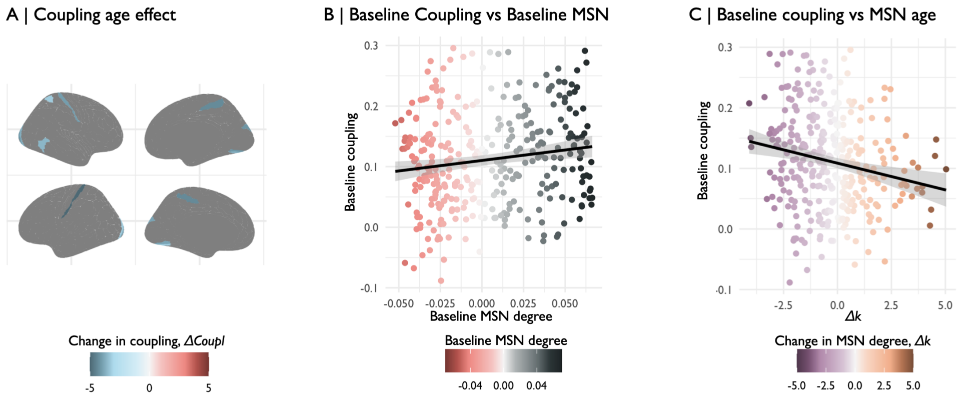

**Fig. 20: Structure-function coupling:** First, we estimated local structure-function coupling as the correlation between a subjects ranked vector of edgewise MSN and FC connectivity values at each region. Then, we estimated the linear effect of age on structure-function using a linear mixed effects model, with a fixed effect of age, sex and site, and a random effect of subject. We derived the baseline structure-function coupling as the predicted coupling at age 14; and the rate of change in coupling as the age effect from this model. (A) We found that 10 regions show significant changes in structure-function coupling after correction for multiple comparisons ($P_{FDR} < 0.05$). (B) Further, we found that the morphometric similarity at baseline (**SI Fig. 21**) was significantly correlated with the baseline structure-function coupling, thus regions with increased morphometric similarity showed increased structure-function coupling. (C) Lastly, we found that there was a significant negative correlation between the age effect on morphometric similarity and the baseline structure-function coupling, meaning that regions that decreased in morphometric similarity tended to have increased coupling at baseline.

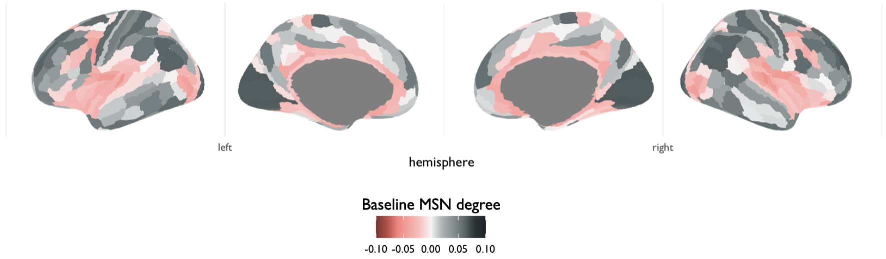

**Fig. 21: Baseline morphometric similarity:** We estimated the linear change with age in regional morphometric similarity degree, $k$, using using linear mixed effects models with a fixed effect of age, sex and site, and a random effect of subject. From this model, we estimated baseline morphometric similarity as the predicted morphometric similarity at age 14. Here, we show the regional baseline morphometric similarity.

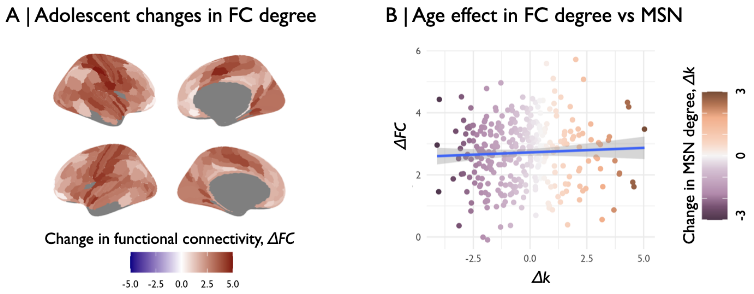

**Fig. 22: Adolescent changes in morphometric similarity are not associated with changes in functional connectivity weighted degree:** Using subjects' functional connectomes, we estimated the adolescent change in weighted degree of functional connectivity (FC) using linear mixed effects models. (A) We find that regional weighted degree of FC increased across the cortex ($t_{age} > 0$). (B) There was no significant association between age-related changes in weighted degree of FC and MSN ($r = 0.05, P_{spin} = 0.4$).

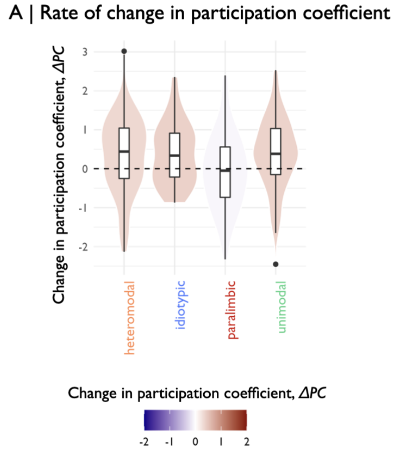

**Fig. 23: Adolescent changes in participation coefficient by Mesulam zone:** We were interested in understanding differences by cytoarchitectonic zones in the rate of change in functional participation coefficient over the course of adolescence. Specifically, we estimated the linear change with age in regional participation coefficient of functional connectivity using using linear mixed effects models with a fixed effect of age, sex and site, and a random effect of subject. From this model, we derived the rate of change in participation coefficient as the $t$-value of the age term. (A) We then averaged the rate of change in functional participation coefficient by Mesulam zone (Mesulam 1998; Mesulam 2000) and find that participation coefficient tended to increase in iso-cortical areas and decrease or remain unchanged in paralimbic areas. See **SI Table 8** for details on pairwise comparisons.

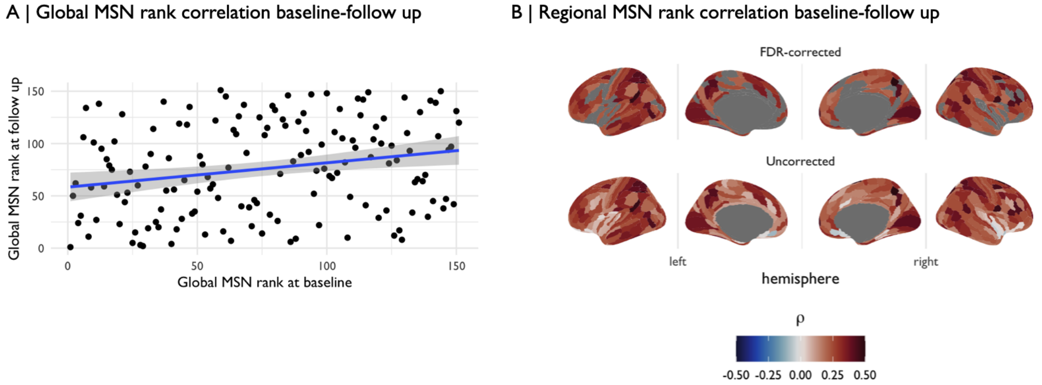

**Fig. 24: Rank-stability of MSN global and regional degree:** We estimated the between-visit rank stability in global and regional MSN degree. To this end, for subjects that had a baseline and one year follow up scan, we ranked subjects by their global and regional MSN degree respectively. We then estimated the Spearman correlation between subjects ranks at baseline and follow. (A) The global rank at baseline compared to follow up was significantly positively correlated ($\rho = 0.23, P < 0.01$). (B) Regionally, we find largely significant positive correlations between subjects' regional MSN degree rank at baseline compared to the follow up scan. The correlation was significant after correction for multiple comparisons for 241 regions ($P_{FDR} < 0.05$).
